# Population Transcriptome and Phenotype Reveal that *Rht-D1b* Contributes a Larger Seedling Root to Modern Bread Wheat

**DOI:** 10.1101/2022.06.02.494553

**Authors:** Xiaoming Wang, Peng Zhao, Xue Shi, Xiaolong Guo, Yuxiu Liu, Wenyang Hou, Mingzhu Cheng, Xueting Liu, Xiangjun Lai, James Simmonds, Wendy Harwood, Junzhe Wang, Zihui Liu, Liuying Huang, Dejun Han, Wanquan Ji, Cristobal Uauy, Jun Xiao, Zhensheng Kang, Shengbao Xu

## Abstract

The Green Revolution (GR) has dramatically increased the yield of modern wheat cultivars; however, whether and how GR reshaped the wheat root system remains largely unknown. Here, a large-scale transcriptomic and phenotypical investigation was performed on seedling roots of 406 worldwide bread wheat accessions, showing differential transcriptomes and phenotypes between landrace and modern cultivars, and the GR allele *Rht-D1b* was the main genetic factor driving this phenotype differential, conferring a significantly larger seedling root to modern cultivars by increasing cell length and root meristem size. In this case, the translational reinitiation of *TaRht-D1* underlies *Rht-D1b*’s genetic effects. In contrast, another GR allele, *Rht-B1b*, has no significant effects on root-related traits, differing from the similar genetic effects of these two GR alleles on reducing plant height. This unexpected effect of *Rht-D1b* on root systems contributes to a substantially larger root-shoot ratio in modern wheat cultivars. These findings reveal previously overlooked benefits of GR alleles in modern wheat cultivars and provide clues for their future application to form a robust seminal root system.

## Introduction

Roots are the fundamental system used for water and nutrient uptake^1^. Bread wheat (*Triticum aestivum* L.) is a hexaploidy comprising A, B, and D subgenomes, and its monocotyledonous root is composed of seminal and adventitious roots^2,3^. The seminal root system is the first to develop, consisting of the primary root and several seminal roots^4,5^. Seminal root development directly influences the formation of the shoot system as well-developed roots support the growth of stems, leaves and other aerial parts, ultimately influencing biomass accumulation and final yield potential^6-9^. Moreover, the well-established seminal root system plays a critical role in improving seedling’s tolerance to abiotic stresses, such as drought and nutrient deficiencies^10,11^. Therefore, the seminal root system is not only pivotal for seedling establishment but also for enhancing stress resistance and ensuring high yield potential.

Over the past century, the breeding selections targeting high-yield have remarkably improved wheat grain yield^12-14^, largely altered genome structures and differentiated modern cultivars (MC) from landraces (LA)^3,15,16^. Significantly, the Green Revolution (GR) of the 1960s dramatically increased grain yields by adopting high-yielding semi-dwarf varieties and increasing nitrogen-based fertiliser using^17,18^. In wheat, the dwarfing was achieved by introducing the *Rht* (Reduced height) alleles *Rht-B1b* and *Rht-D1b* (also named *Rht1* and *Rht2*), which induce premature stop codons in the N-terminal coding region of DELLA, a key functional repressor of gibberellin (GA) signalling, conferring dominant GA-insensitive dwarfism^19,20^. After six decades of practice, these two alleles are now present in >70% of current commercial wheat cultivars worldwide^21,22^. Notably, the shorter coleoptiles associated with the wheat GR alleles limit sowing into deep moisture, making the deeper and larger seminal root system more crucial for the GR varieties^7^. However, the effects of wheat GR alleles on the seminal root system have not drawn consistent conclusions due to the limited measured traits and the difference in used genetic backgrounds and cultivation substrates^23,24^.

With increased sequencing accuracy, large-scale population transcriptome sequencing is becoming an important tool for population genetic analysis^25,26^. It determines the transcriptional levels of each gene among the accessions that cannot be captured at the DNA sequence level^27^, thus building a bridge from sequence polymorphisms to phenotype variations and providing critical clues for the prioritization of causal genes and the underlying mechanism investigations of identified functional alleles^28,29^.

Here, we analysed the root system’s transcriptomes and phenotypes of 406 worldwide bread wheat accessions, revealing that the breeding selection substantially altered root transcriptomes and development. GWAS and integrative transcriptome analysis demonstrated the GR allele *Rht-D1b*, differentiated from the *Rht-B1b*, to be the main genetic factor contributing to larger seminal root systems of GR varieties. Cytological analysis showed that *Rht-D1b* enlarged root meristem size and increased cell length and width in the mature region, opposite to its well-known dwarfing effects on the plant height, thereby significantly increasing the root-shoot ratio of GR varieties. These results further explain how GR alleles contribute to wheat production at the seedling stage and provide new insights into their application in future seminal root improvement.

## Results

### A large-scale transcriptome sequencing reveals the gene composition and variations in wheat seminal root

A large-scale RNA-seq was performed on 406 worldwide bread wheat accessions with the seedling root samples at 14 days after germination (DAGs), producing 15.98 billion high-quality reads and identifying 1,232,312 single nucleotide polymorphisms (SNPs) with an average density of 10 SNPs per gene (Fig. 1a, Fig. S1 and Table S1). The accuracy of the identified SNP was validated as > 99% with wheat SNP microarray evaluation (Methods and Table S2). About half of the SNPs (48.43%) were located in CDS (coding DNA sequence) regions (Fig. 1b), consistent with the feature of transcriptome-derived SNPs.

**Figure 1.**
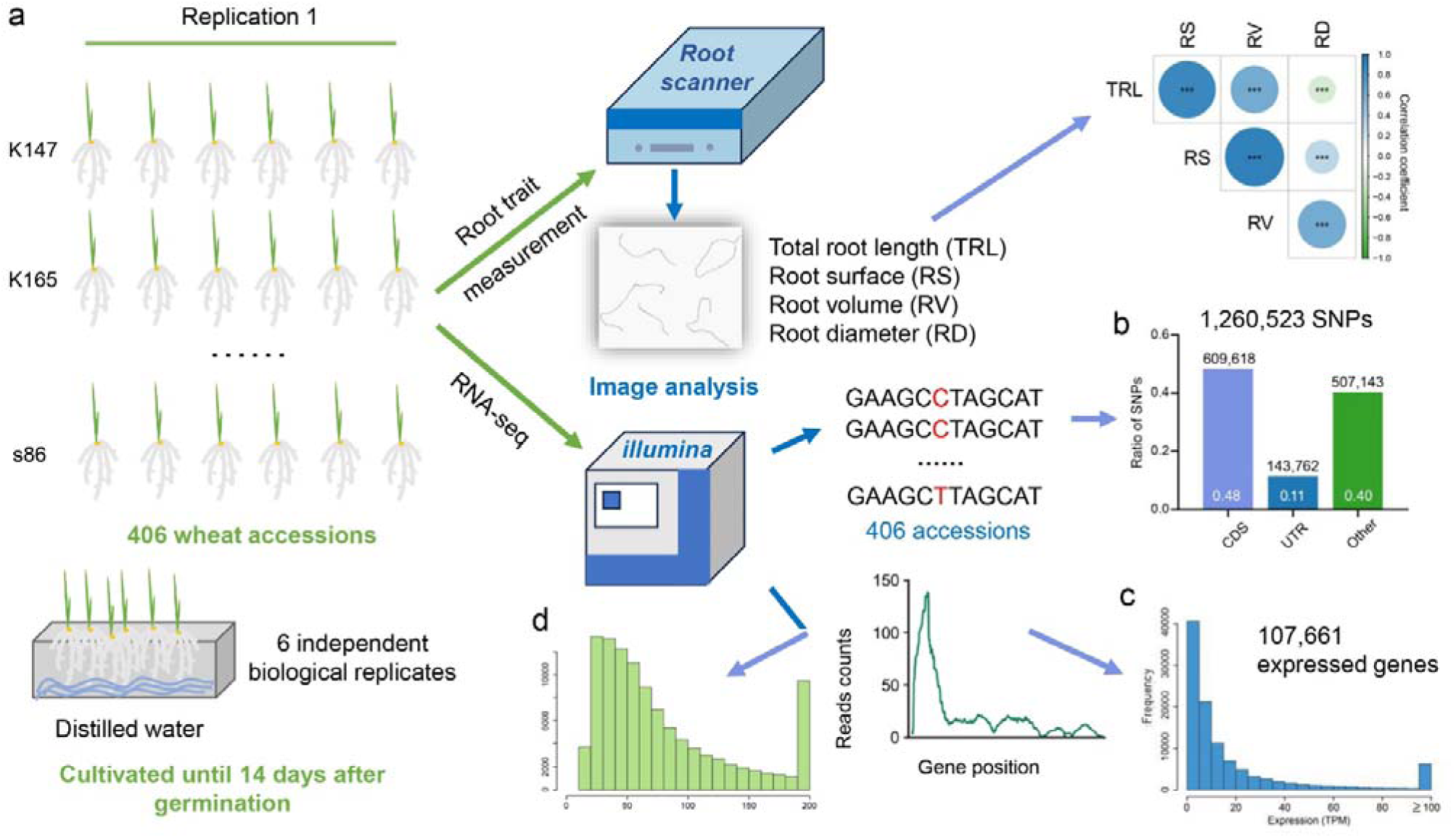
Population transcriptome in wheat root. (a) Overview of the phenotyping and genotyping workflows. (b) The distribution of the identified SNPs in gene regions. (c) The transcripts per million (TPM) distribution of expressed genes. (d) The coefficient of variation of expressed genes.

With this population transcriptomes, 107,661 genes were identified as expressed in root with transcripts per million (TPM) > 0.5 in at least 20 accessions, including 39,243 low confidence (LC) genes, providing further transcriptional evidence for these gene annotations. The expression levels of these genes ranged from 0.5 to 21,144.5 TPM with the median as 2.01 (Fig. 1c). The enriched gene ontology (GO) terms for the highly expressed genes include “response to water deprivation” and “water transport” pathways (Fig. S2a and Table S3), consistent with the fundamental functions of root systems. Interestingly, these two GO terms were also enriched in the genes with highly varied expression levels across the accessions (Fig. 1d, Fig. S2b and Table S4), implying varied root functions in water use efficiency among accessions. The identified SNPs and determined TPM values can be accessed and downloaded from (http://resource.iwheat.net/WGPD/).

### Modern wheat breeding substantially changed the seminal root transcriptome and phenotype

Phylogenetic and population structure analyses clearly separated 190 modern cultivars (MC, released after GR) and 87 landraces (LA) from the used 406 wheat accessions at the DNA sequence level (Fig. 2a and 2b). Interestingly, the root transcriptomes of MC and LA accessions were also generally separated in the PCA (Principal Component Analysis) plot (Fig. 2c). Furthermore, after population structure correction, 8,179 differentially expressed genes (DEGs) were identified between LA and MC groups (FDR < 0.05 and fold change ≥ 1.2; Table S5), including 869 MC-private (expressed in >= 10% MC accessions while in < 5% LA accessions) and 853 LA-private (expressed in >= 10% LA accessions while in < 5% MC accessions) expression genes. These pieces of evidence demonstrated the root transcriptomes were significantly changed during modern wheat breeding selection. Gene Ontology (GO) enrichment analysis of the DEGs indicated that the plant development, phytohormones signalling transductions, environmental stress response and many catabolic processes were significantly changed between LA and MC groups (Fig. 2d and Table S6). Notably, the homologs of 39 DEGs in other plants were identified as the key genes involved in root development (Table S7), such as *AtRCD1*^30^, *AtARR10^31,32^*, *AtEXPB2*^33^, *OsSHB*^34^, *OsAUX3*^35^, *OsPIN1/3/7*^36^ and *OsERF2^37^*.

**Figure 2.**
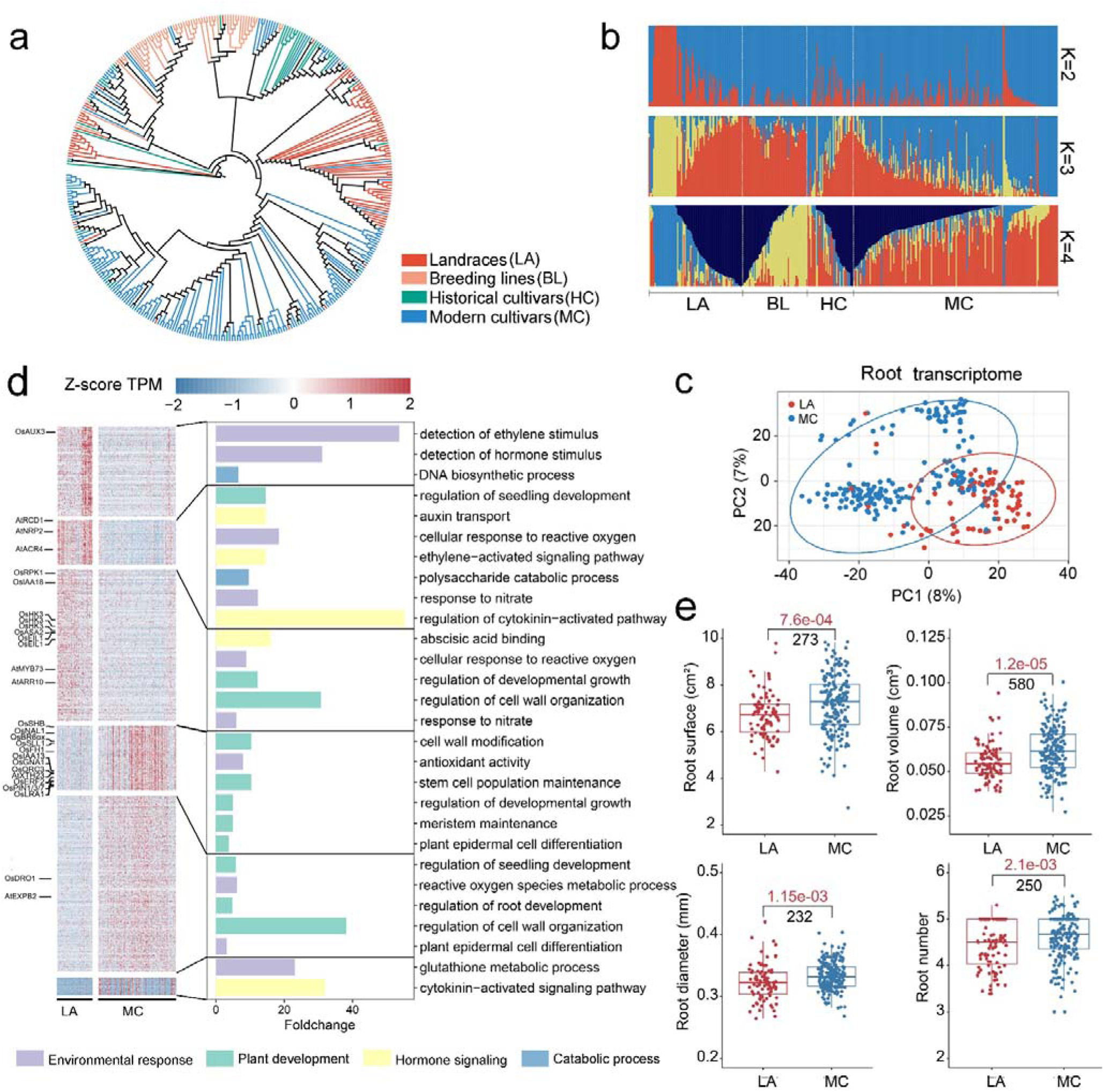
The changes of root transcriptome and root development in modern wheat cultivars. (a) The phylogenetic tree of 406 wheat accessions, constructed with the identified SNPs. The colours represent different groups. LA: landraces (red); HC: historical cultivars (green, released before Green Revolution); MC: modern cultivars (blue, released after Green Revolution); BL: breeding lines (orange). (b) Population structure of 406 wheat accessions with K (hypothetical subpopulations) = 2, 3, and 4. Each vertical line represents an accession. (c) PCA plots of the first two components calculated using the root transcriptomes of LA and MC accessions. (d) Gene expression and GO enrichment of the differentially expressed genes (DEGs) between LA and MC groups. Each column represents an accession. The DEGs whose homologs in model plants were reportedly involved in root development were identified, and the gene names in model plants were labelled on the left. (e) The comparisons of root-related traits between LA (n= 87) and MC (n= 190). Data are means ± SD. Red numbers indicate the *P-*value from the Student’s *t*-tests. Black numbers indicate the observed number with significant differences at *P*< 0.01 in the 1,000 permutation tests.

At last, the system measurements of root-related traits showed the root surface (RS), root volume (RV), root diameter (RD) and root number were significantly increased (*P* < 0.01) in the MC group compared with that in the LA group with both original phenotype values (Fig. 2e) and the corrected values by excluding the effects of grain weight^38^, providing experimental evidence that the modern wheat breeding substantially changed seminal root development.

### Dissection of the relationship between the changed root transcriptomes and phenotypes

To investigate the relationship between the changed transcriptomes and phenotypes, we first performed Transcriptome-Wide Association Studies (TWAS) with the mixed linear model (MLM) in which the population structure and family relatedness were corrected, identifying 16,717 genes whose expression levels were significantly associated with at least one root-related trait at the genome-wide level (FDR < 0.05, Fig. 3a-b and Table S8). Among them, the homologs of 73 genes were reportedly involved in root development in the model plants, supporting the present analytic strategy and statistical power of the MLM analysis (Fig. 3a and Table S9). This gene set includes the following: *TraesCS2A02G192600*, *TraesCS2B02G214600,* and *TraesCS2D02G194800* are homologoues of the *Arabidopsis SHORTROOT* (*SHR*), which positively regulate the development of endodermis and cortical tissue^39^; *TraesCS6A02G308600* is homologoue of the *Arabidopsis PIN-FORMED 1* (*PIN1*) which is auxin efflux carrier and essential for establishing endogenous auxin gradients and root development^36^.

**Figure 3.**
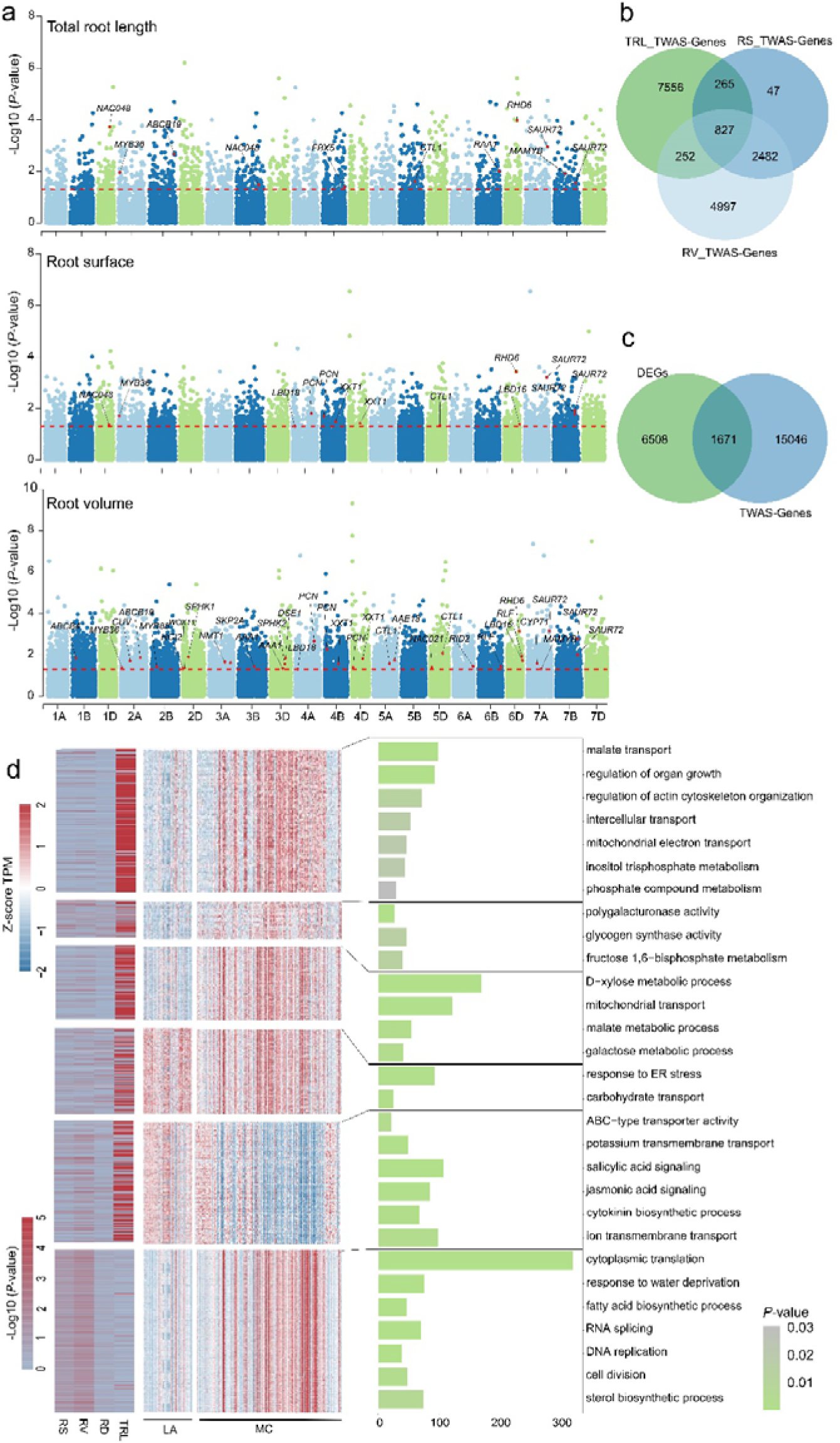
Investigation of the relationship between transcriptomes and phenotypes with TWAS. (a) The results of TWAS analysis on root-related traits. The red line in each plot represents the significance of FDR < 0.05. (b) The overlapped genes are associated with different root traits. “TRL”, “RS” and “RV” represent the total root length, root surface and root volume, respectively. (c) The overlapped set between genes significantly associated with TWAS analysis and differentially expressed genes (DEGs) between LA and MC groups. (d) Expression patterns, significant levels in TWAS and GO enrichments of the overlapped genes in (c). The left section represents the significance of TWAS analysis. In the middle, each column represents an accession.

For the above-mentioned 8,179 DEGs between MC and LA groups, 1,671 of which were significant in TWAS analysis, which may be responsible for the differential phenotypes in these two groups (Fig. 3c). Using the k-means clustering algorithm, we classified the 1,671 overlapped genes into six clusters (Fig. 3d). The cluster 1, 2 and 3 generally had similar patterns that these genes were up-regulated in the MC group, and were enriched in the GO terms related with biological macromolecule metabolisms (Fig 3d and Table S10). Interestingly, clusters 5 and 6 had opposite expression patterns, showing that the genes in cluster 5 were down-regulated while those in cluster 6 were up-regulated in the MC group. Cluster 5 was enriched in the GO terms of ion transmembrane transport and phytohormone signalling (Salicylic acid, Jasmonic acid and Cytokinin), and cluster 6 was enriched in the transcription, translation and cell division processes. These results suggested that the biological macromolecule synthesis and the basic metabolism processes were accelerated by the breeding improvement, which may account for GR varieties’ larger seminal root systems.

### GR allele *Rht-D1b* contributes larger seminal root systems to modern cultivars

To identify the genomic regions responsible for differentiating root transcriptomes and phenotypes of the MC group from the LA group, Genome-Wide Association Studies (GWAS) were performed, which detected 381 pairs of associations between SNPs and root-related traits (Fig. 4a and Table S11). Then, these significantly associated SNPs were aligned to our previously identified selection sweeps during wheat breeding^38^, finding 33 SNPs located in the selection sweeps, which were named MC-LA-SNPs (Table S11). To filter out the MC-LA-SNPs playing major roles, the associations between the SNPs and the expression levels of each root-expressed gene were constructed with GWAS using the gene expression level as phenotype, revealing 9,681,969 pairs of associations at the genome-wide level (http://resource.iwheat.net/WGPD/). In these results, we found 1,265 pairs of associations between the MC-LA-SNPs and the root-expressed genes. Further, we focused on 452 pairs of the associations in which the regulated genes were significantly associated with root-related traits in the above TWAS analysis (Fig. 4b-c, Fig. S3 and Table S12). These MC-LA-SNPs and associated genes may play important roles in differentiating the root systems of the MC group from the LA group by connecting the DNA variants, transcriptome alteration and phenotype changing.

**Figure 4.**
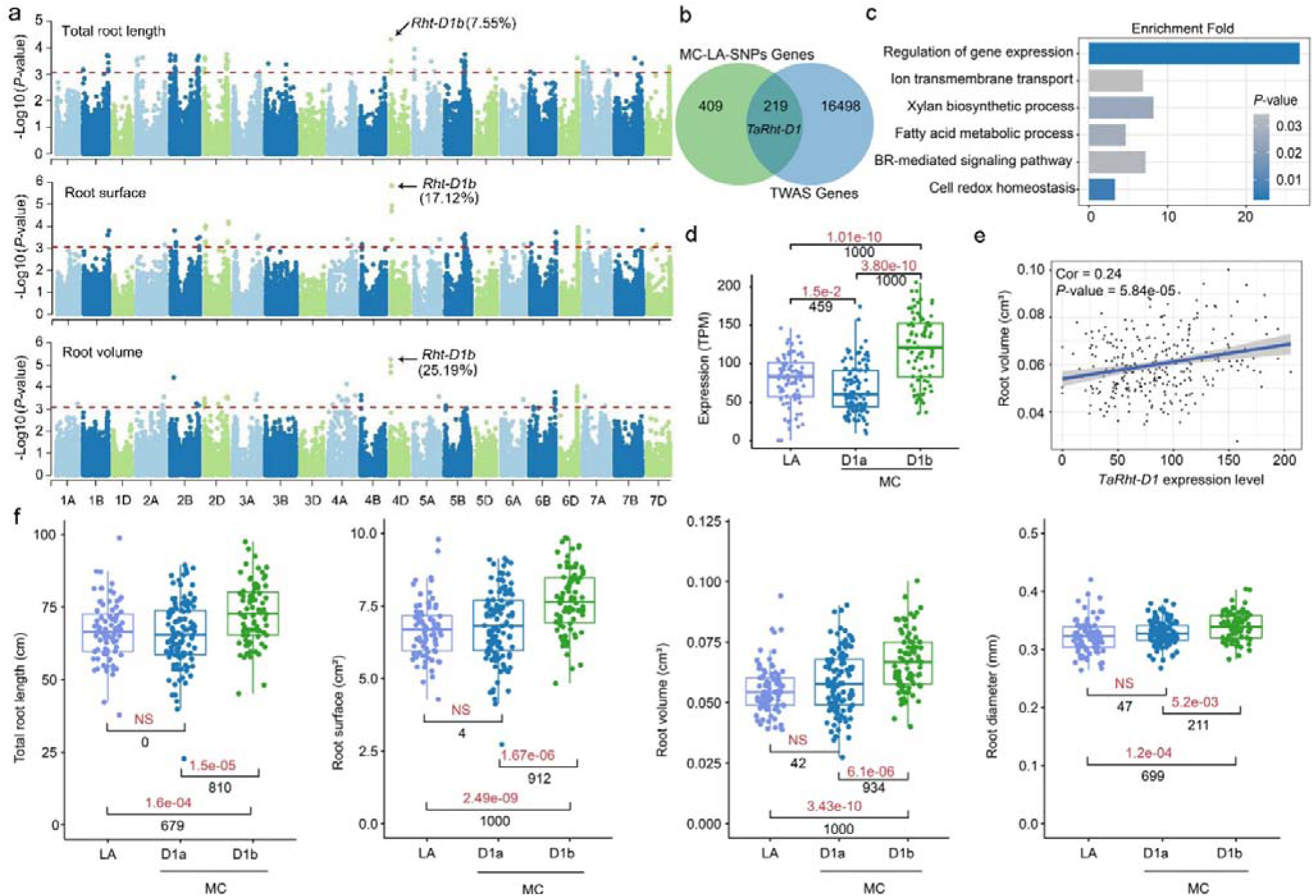
The *Rht-D1b* contributes a larger seedling root system to modern wheat cultivars. (a) The results of GWAS analysis on root-related traits. The red line in each plot represents the significant *P-*value threshold calculated with Bonferroni-corrected threshold probability. The numbers indicate the phenotypic variance explained by *Rht-D1b*. (b) The overlapped set between genes whose expression levels were significantly associated with MC-LA-SNPs and genes significantly associated with TWAS analysis. (c) The GO enrichment of the overlapped genes in (b). (d) The expression levels of *TaRht-D1*. (e) Correlations between root volume and the expression level of *TaRht-D1*in the whole panel. (f) The comparisons of root-related traits between LA (n = 87) and MC (n = 190) groups. (d, f) The MC accessions were divided into *D1a* (n = 110) and *D1b* (n = 80) genotypes based on the GR allele *Rht-D1b*. Red numbers indicate the *P-*values from the Student’s *t*-test, and “NS” represents insignificance at the *P*-value > 0.01 level. Black numbers indicate the observed number with significant differences at *P*-value < 0.01 or insignificant differences at *P*-value > 0.01 level (for the “NS”) in the 1,000 permutation tests.

Among the above-established associations, we were interested in the association between MC-LA-SNP chr4D&18781242 and *TraesCS4D02G040400*. The chr4D&1878124 2corresponds to the well-known GR allele *Rht-D1a* (75.86%) and *Rht-D1b* (24.14%), which was significantly associated with total root length (TRL), RS, and RV and explained 7.55%, 17.1% and 25.2% phenotype variations in the GWAS, respectively (Fig. 4a). The *Rht-D1b* genotypes had significantly larger TRL, RS, RV and RD than the *Rht-D1a* genotypes (Fig. S5a). The *TraesCS4D02G040400* was annotated as *TaRht-D1*, whose expression levels were significant with RS, RV and RD (Fig. 4d-e and Fig. S4). Considering the effect of kernel weight on seedling root development, we corrected the root phenotype values to exclude its effect with the general linear model. The corrected traits were significantly larger in *Rht-D1b* genotypes (Fig. S5b), and this SNP was still significantly associated with the corrected traits in the GWAS analysis (Fig. S6 and Table S13), and the expression levels of *TaRht-D1* were also significantly associated with the corrected traits in the TWAS (Fig. S4b). Significantly, TRL, RS, RV and RD no longer showed the difference between the LA and MC groups if the *Rht-D1b* genotypes were removed from the MC group, strongly supporting that the *Rht-D1b* is the main candidate that enlarged the seminal root system of MC accessions (Fig. 4f and Fig. S7).

Then, to investigate whether the larger root systems of *Rht-D1b* genotypes resulted from earlier germination, we performed systematic and continuous investigations on the root development (Fig. S8). At four DAGs, the primary root length and TRL of *Rht-D1b* genotypes were significantly shorter than that of *Rht-D1a* genotypes. At seven DAGs, larger RV and RD began to be observed in the *Rht-D1b* genotypes. The TRL and RS of *Rht-D1b* genotypes were not significantly larger until 14 DAGs (Fig. S8). These results exclude the possibility that the large roots of *Rht-D1b* genotypes were due to earlier germination.

It is intriguing that another wheat GR allele, *Rht-B1b*, from *TaRht-B1* located on chromosome 4B, was not associated with any root-related traits in GWAS analysis; the investigated traits did not show a significant difference between *Rht-B1a* and *Rht-B1b* genotypes, and the *TaRht-B1* were also not significant in TWAS analysis (Fig. S9-10), although the *TaRht-B1* and *TaRht-D1* were homoeologs and the positions of *Rht-B1b* and *Rht-D1b* were much closer^40^. To experimentally validate the genetic efforts of *Rht-D1b* and the differential effects between *Rht-D1b* and *Rht-B1b*, the Near Iso-genic Lines (NILs), introducing the *Rht-D1b* and *Rht-B1b* into wheat cv. Paragon (*Rht-B1a* and *Rht-D1a* background), respectively, was constructed and investigated. The Paragon*^Rht-B1aRht-D1b^*(Paragon*^B1aD1b^*) showed larger root systems than Paragon*^Rht-B1aRht-D1a^*(Paragon*^B1aD1a^*). However, the root system of Paragon*^Rht-B1bRht-D1a^*(Paragon*^B1bD1a^*) was comparable with Paragon*^B1aD1a^*(Fig. 5a-b). Notably, consistent results were observed by growing the NILs in other substrates and fields, demonstrating the reliability of the *Rht-D1b*’s effects (Fig. 5c-d and Fig. S11).

**Figure. 5.**
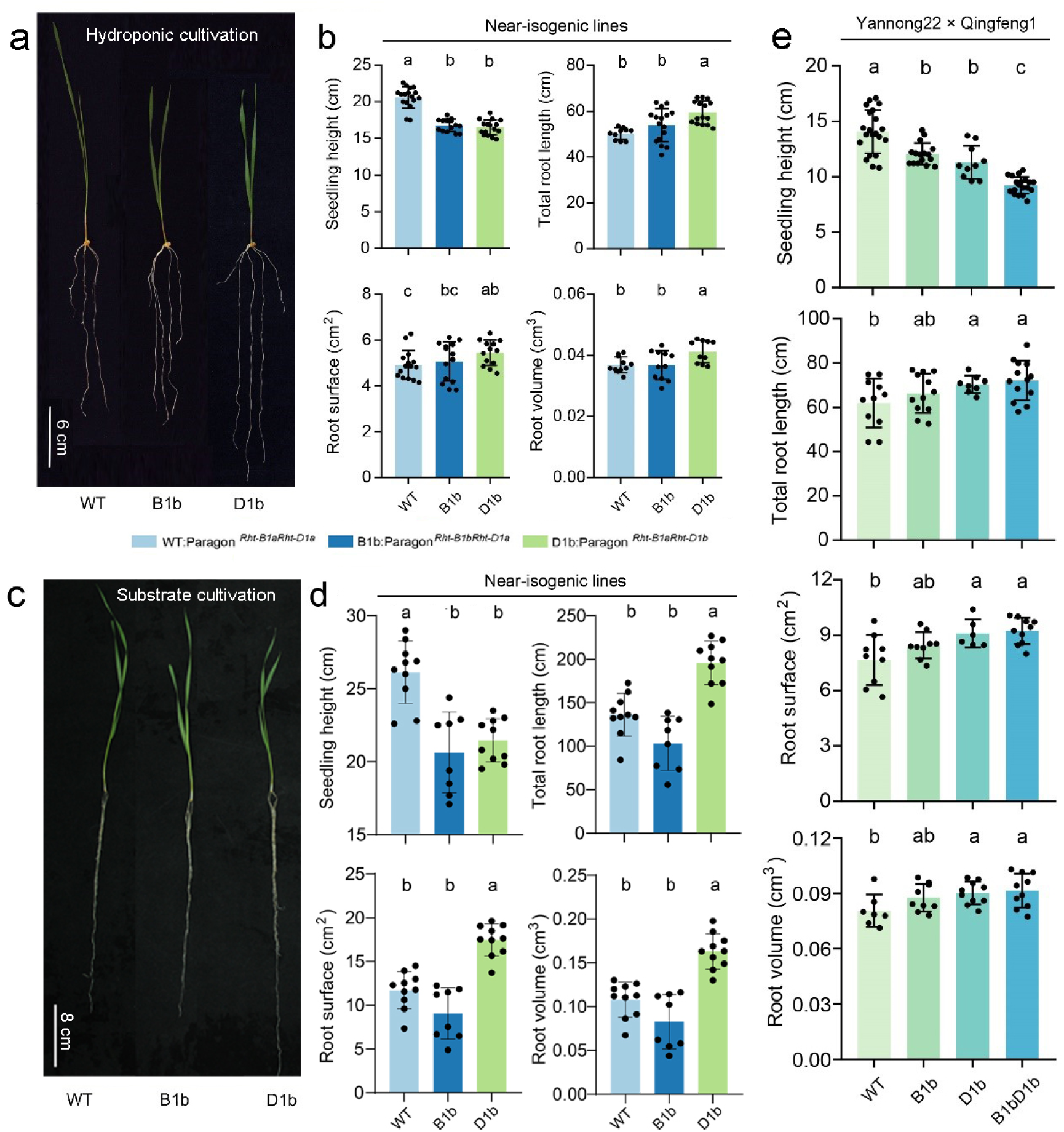
The genetic effects of *Rht-B1b* and *Rht-D1b* on root development. (a-d) Phenotype and statistic data of the Near-isogenic lines cultivated with hydroponic (a and b) and substrates (c and d). (e) Comparison of the root-related traits among different genotypes in the F_2:3_ segregation population. The parents are Yannong22 (*Rht-B1bRht-D1a*) and Qingfeng1 (*Rht-B1aRht-D1b*). Data are means ± SD, and different letters on the top indicate significant differences (*P* < 0.05) calculated with multiple comparisons by the least significant difference (LSD) test.

Next, three F_2_ segregation populations were constructed by hybridising the parent accessions harbouring *Rht-B1aRht-D1b* and *Rht-B1bRht-D1a* alleles further to validate the above results in different genetic backgrounds. Consistent with the results from the NIL lines, the root-related traits showed significant differences between *Rht-B1aRht-D1b* and *Rht-B1aRht-D1a* genotypes in all of the three populations (Fig. 5e and Fig. S12). However, the differences between *Rht-B1bRht-D1a* and *Rht-B1aRht-D1a* genotypes were insignificant, although the *Rht-B1bRht-D1a* genotypes showed larger root systems. These results provided genetic evidence for the function of *Rht-D1b* in enlarging root systems of wheat seedlings and the functional differentiation between the two wheat GR alleles in root development.

### Transgenic validation of the genetic effects of *Rht-D1b*

To validate the function of *Rht-D1b* in regulating root development, the full CDS of *TaRht-B1* and *TaRht-D1* containing the *Rht-B1b* and *Rht-D1b* alleles (F-*Rht-B1b-OE* and F-*Rht-D1b-OE*, Fig. 6a), respectively, driven by their native promoters were cloned into wheat cv. Fielder (*Rht-B1bRht-D1a* background). The transgenic lines showed significantly higher expression levels of *TaRht-B1* and *TaRht-D1* and reduced seedling height (Fig. 6b-c and Fig. S13), consistent with their well-known dwarfing functions. For the root-related traits, the two F-*Rht-D1b-OE* lines showed larger root systems than the wild type (Fig. 6c). In contrast, the F-*Rht-B1b-OE* lines didn’t show significant increases in root-related traits, although the *TaRht-B1* transformations contribute a similar increase in the transcription levels with that of *TaRht-D1* (Fig. 6c).

**Figure 6.**
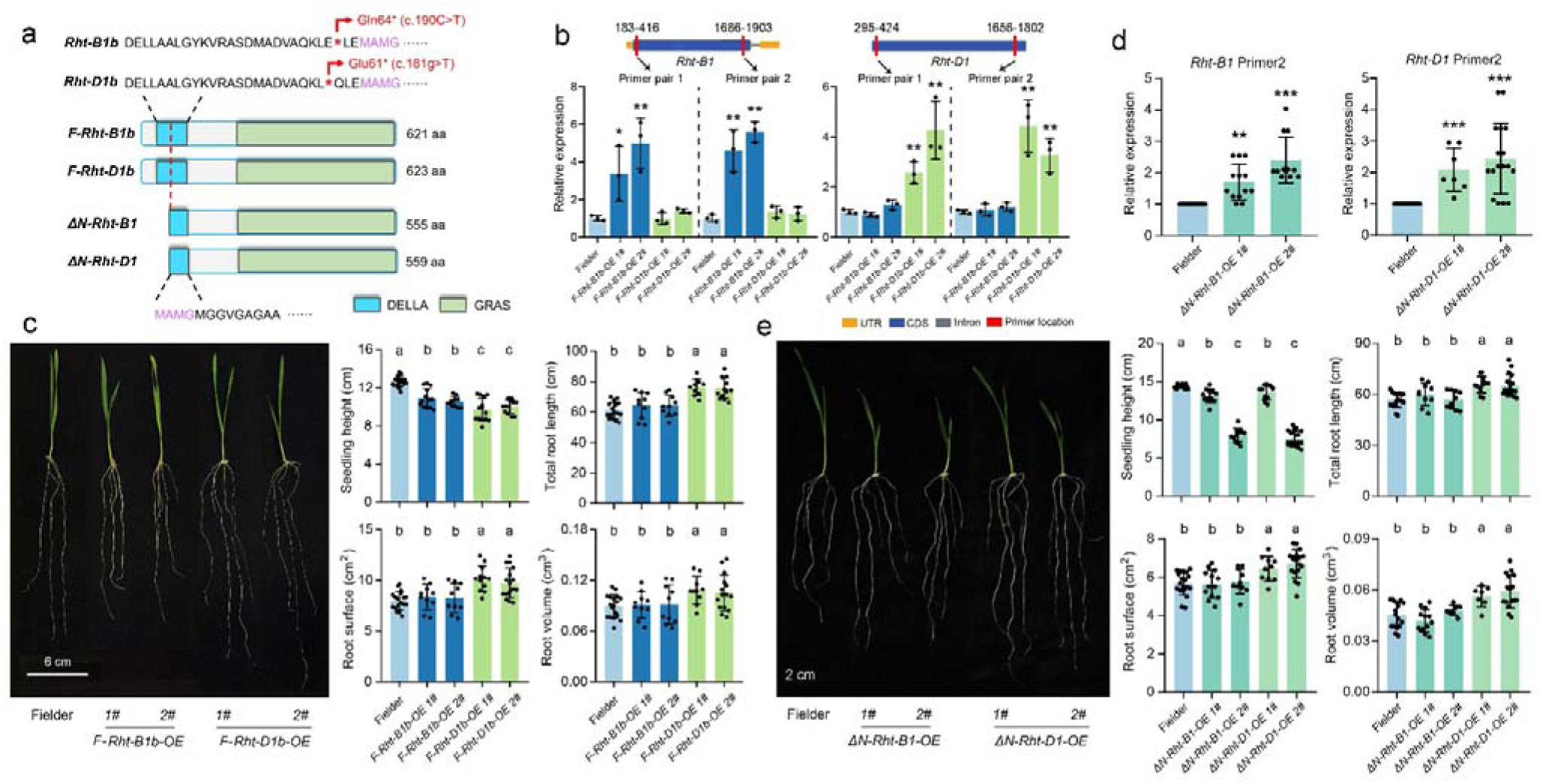
Validation of the effects of *Rht-B1b* and *Rht-D1b* on root development with transgenic assays. (a) The model of CDS sequences used for transgenic assays. F-*Rht-B1b* and F-*Rht-D1b* represent the full CDS of *Rht-B1* and *Rht-D1*, containing the GR alleles; ΔN-*Rht-B1* and ΔN-*Rht-D1* represent the N-terminal truncated CDS of *Rht-B1* and *Rht-D1* beginning from the first start codon after GR alleles. All the CDS were transformed into wheat cv. Fielder driven by their native promoters. (b) The expression levels of *Rht-B1* and *Rht-D1* in the F-*Rht-B1b* and F-*Rht-D1b* transgenic lines. Three independent plants were quantified per line. Two pairs of primers were used for *Rht-B1* and *Rht-D1*, respectively, in the qRT-PCR analysis. “*” and “**” indicate significant differences (Student’s *t*-test *P* < 0.05 and 0.01) between the wild-type and transgenic lines. Fielder: wild-type; *Rht-B1b-OE 1/2#*, F-*Rht-B1b* transgenic lines; *Rht-D1b-OE 1/2 #*: F-*Rht-D1b* transgenic lines. (c) Root-related phenotypes and statistical data of the wild-type, F-*Rht-B1b* and F-*Rht-D1b* transgenic lines. Data are means ± SD, and different letters (a, b, and c) on the top indicate significant differences (*P* < 0.05) between different genotypes with multiple comparisons by least significant difference (LSD) test. (d) The expression levels of *Rht-B1* and *Rht-D1* in the ΔN-*Rht-B1* and ΔN-*Rht-D1* transgenic lines. Fielder: wild-type; Δ*N-Rht-B1-OE 1/2#*, ΔN-*Rht-B1* transgenic lines; Δ*N-Rht-D1-OE 1/2 #*: ΔN-*Rht-D1* transgenic lines. (e) Root-related phenotypes and statistical data of the wild-type, ΔN-*Rht-B1* and ΔN-*Rht-D1* transgenic lines.

The *Rht-B1b* and *Rht-D1b* induce premature stop codons in the N-terminal coding region of DELLA, a key repressor of the GA signalling. They might be expected to prevent translation into a functional protein. However, *Rht-B1b* and *Rht-D1b* are considered to be gain-of-function alleles that confer dominant GA-insensitive dwarfism^41^. The translational reinitiation (TR), a process where ribosomes bypass premature stop codons to resume protein synthesis downstream, generating N-terminal truncated proteins lacking the DELLA domain, was proposed to account for this phenomenon, and the molecular evidence was reported recently in spikes and nodes^18,19,41^. To test whether the TR and N-terminal truncated proteins were also involved in the observed functions of *Rht-D1b* in regulating root development, an N-terminal truncated CDS of *TaRht-B1* and *TaRht-D1*, beginning from the first methionine start codon following the mutant stop codons were cloned and transformed into Fielder under their native promoters, respectively (ΔN-*Rht-B1-OE* and ΔN-*Rht-D1-OE*, Fig. 6a). Both ΔN-*Rht-B1-OE* and ΔN-*Rht-D1-OE* showed significantly higher expression levels of the target CDS fragment and much more reduced seedling height (Fig. 6d-e and Fig. S13). For the root-related traits, like the F-*Rht-D1b-OE* lines, the two ΔN-*Rht-D1-OE* lines showed significantly increased TRL, RS and RV compared to the wildtype, demonstrating the N-terminal truncated CDS of *Rht-D1* did have functions in regulating root development and implying TR underly the mechanism of *Rht-D1b*’s genetic effects (Fig. 6e). In contrast, the ΔN-*Rht-B1-OE* lines showed comparable root-related traits with the wild type although they showed the expected dwarfing plant height. These results demonstrate the function of *Rht-D1b* in enlarging the root system and provide further evidence for the differential effects of the two wheat GR alleles.

### Increased cell length and width underlie the larger seminal root systems of *Rht-D1b* genotypes

To clarify the cytological foundations of *Rht-D1b* in regulating root development, the root meristems of NIL lines and transgenic lines were measured. The lines containing the *Rht-D1b* allele (Paragon*^Rht-B1aRht-D1b^* and F-*Rht-D1b*-OE) showed significantly increased meristem length and width compared to the wild type (Paragon*^Rht-B1aRht-D1a^* and Fielder) while the Paragon*^Rht-B1bRht-D1a^*showed comparable meristem size with the wild type (Fig. 7a and Fig. S14). In detail, the cell width at both distal and proximal meristematic regions increased in the *Rht-D1b* lines, while the transverse cell numbers were comparable (Fig. 7a). The longitudinal cell numbers of *Rht-D1b* lines increased, but the cell length was similar to the other two types of lines (Fig. 7a). For the cells in the mature region, both the cell length and width significantly increased in the *Rht-D1b* lines compared to those in the *Rht-B1b* lines and wild type. However, the ratio of length/width of *Rht-D1b* lines was similar to the other two types of lines (Fig. 7b).

**Figure 7.**
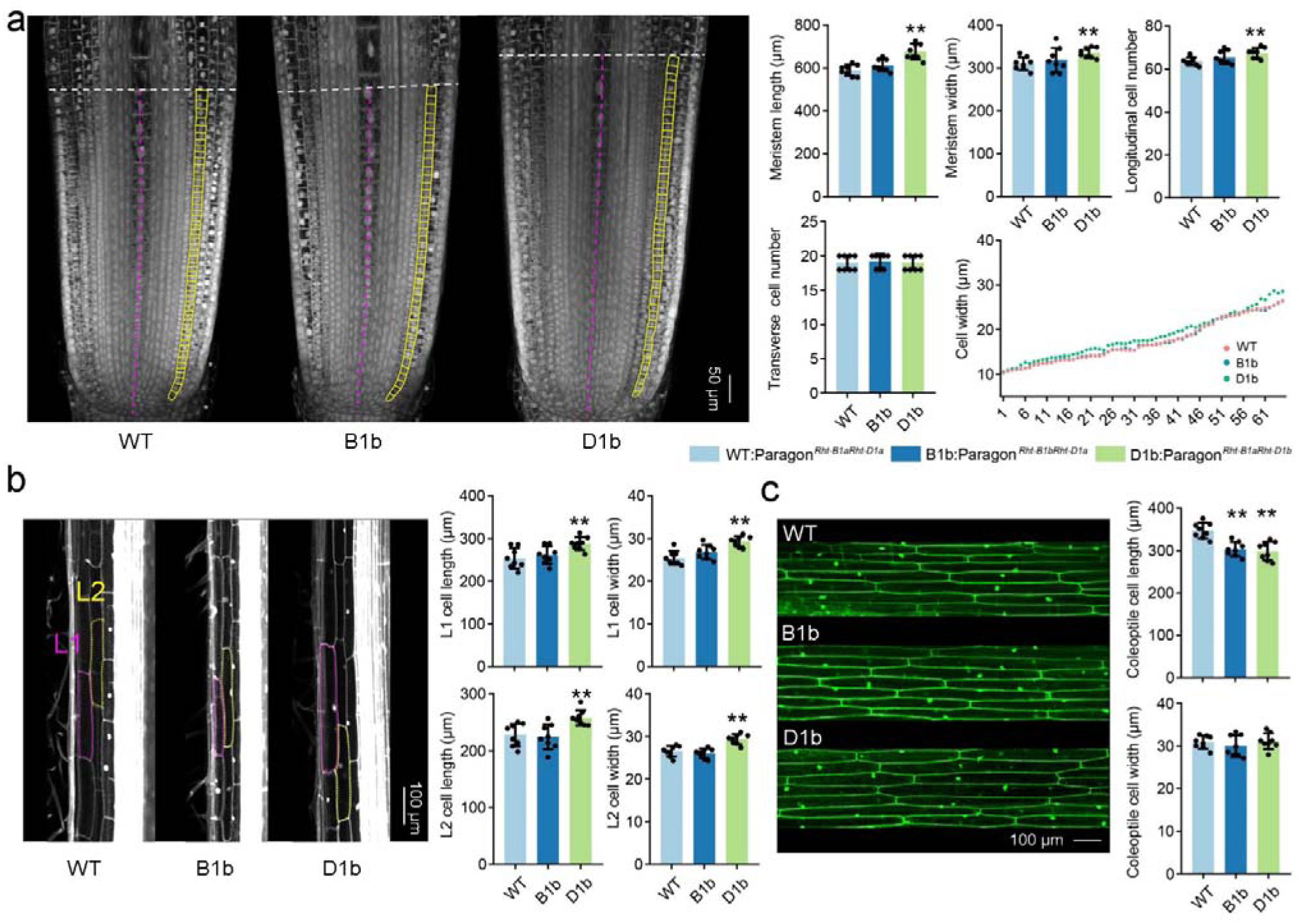
*Rht-D1b* increased the cell length and width in the root. (a) Cytology of root meristem zone of the near-isogenic lines. “Meristem length” and “Meristem width” were indicated by purple and white dotted lines, respectively. The longitudinal cell length and number were measured from the cell layer marked with yellow. (b) Cytology of root maturation zone. “L1” and “L2” represent different cell layers indicated by the purple and yellow dotted lines. (c) Cytology of the coleoptile. At least seven samples were observed for each near-isogenic line, and at least 30 clear cells in each sample were measured with ImageJ software (v1.48). “*” and “**” indicate significant differences (Student’s *t*-test *P* < 0.05 and 0.01) between the compared lines.

In contrast, the *Rht-B1b* and *Rht-D1b* containing lines showed significantly decreased coleoptile cell length while similar cell width to the wild type (Fig. 7c and Fig. S14), consistent with their well-known dwarf phenotype. This highlights the opposite regulation effects of *Rht-D1b* between above-ground and below-ground tissues.

### *Rht-D1b* regulates distinct genes compared with *Rht-B1b*

To understand the molecular mechanisms underlying the functional difference between the *Rht-D1b* and *Rht-B1b* in regulating root development, firstly, the sensitivity of NILs to exogenous GA and PAC (Gibberellin biosynthesis inhibitor paclobutrazol) was evaluated, showing the Paragon*^B1aD1b^*generally was less sensitive than Paragon*^B1bD1a^* regarding the root-related traits, although they were both insensitive to the treatments regarding seeding height (Fig. S15). Then, the genes that respond to the introductions of *Rht-D1b* and *Rht-B1b* were identified by comparing the RNA-seq data of Paragon*^B1aD1b^* and Paragon*^B1bD1a^* to that of the Paragon*^B1aDa^*, respectively. The two gene lists were largely different, making the Paragon*^B1aD1b^* farther from Paragon*^B1aDa^* in the PCA plot (Fig. 8a-b and Table S14-15). Next, we further analyzed whether the private response genes of these two alleles were homoeologous copies located on different subgenomes and found 98.89% and 97.55% of *Rht-D1b*’s and *Rht-B1b*’s private response genes do not have homoeologous in the other gene list, respectively, further supporting these two alleles regulated different genes. GO enrichment showed that the *Rht-D1b*’s private response genes were mainly involved in cell division, redox homeostasis, auxin-stimulus response and phloem development processes. In contrast, the *Rht-B1b*’s private response genes were involved in response to ROS and response to GA stimulus (Fig. 8c and Table S16).

**Figure 8.**
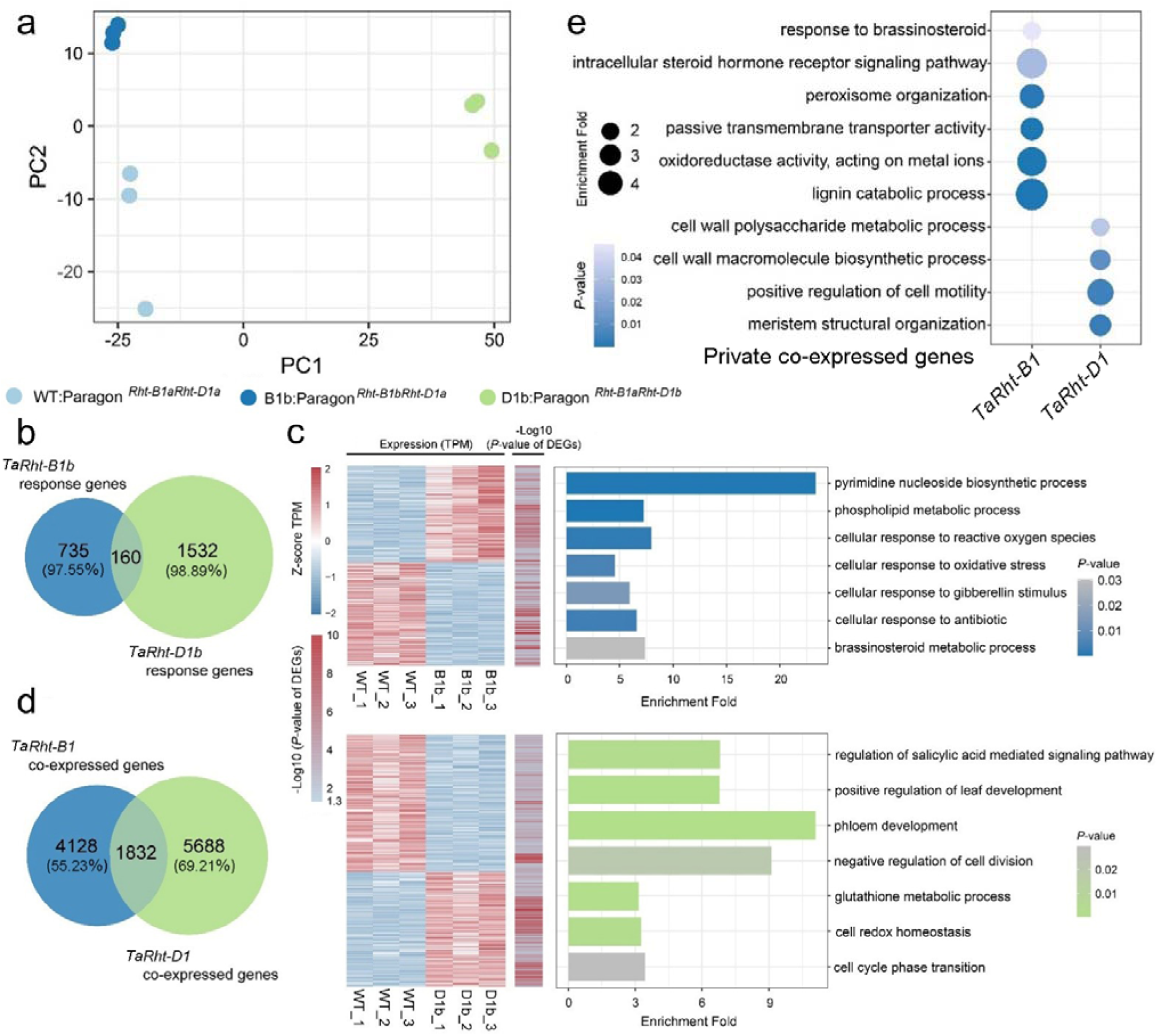
*Rht-D1b* regulates different genes compared with *Rht-B1b* in the root. (a) PCA plots of the first two components calculated with the root transcriptomes of near-isogenic lines. (b) The comparison of genes response to the introduction of *Rht-D1b* or *Rht-B1b*. The numbers in parentheses present the genes that do not have homoeologous copies in the other gene list. (c) The heatmap showed the TPM in the three replications and the significance of the differential expression. The right part shows the GO enrichment results of the *Rht-D1b* and *Rht-B1b* private response genes. (d) The comparison of genes co-expressed with *Rht-D1* or *Rht-B1* in the population transcriptomes. The numbers in parentheses present the genes that do not have homoeologous copies in the other gene list. (e) GO enrichment of the *Rht-D1* and *Rht-B1* private co-expressed genes.

Considering the significant correlation between the expression levels of *TaRht-D1* and root-related traits, the genes co-expressed with *TaRht-D1* and *TaRht-B1* were identified, respectively, in the *Rht-B1aRht-Da* genotypes with our previously developed method^42^. The two co-expressed gene lists were also largely different (Fig. 8d and Table S17), and 69.21% and 55.23% of the genes co-expressed with *TaRht-D1* and *TaRht-B1* do not have homoeolog copies in the other gene list. Besides, the private co-expressed genes with *TaRht-D1* and *TaRht-B1* showed largely different enriched GO terms, such as the root meristem growth, cell cycle and number, cell wall biogenesis and cytokinetic process were enriched for the *TaRht-D1*’s private co-expressed genes. In contrast, brassinosteroids homeostasis, auxin transmembrane transporter activity, and oxidoreductase complex were enriched for the *TaRht-B1*’s private co-expressed genes (Fig.8e and Table S18). These pieces of evidence suggest that *Rht-D1b* and *Rht-B1b* regulate distinct genes, and *Rht-D1b* and *TaRht-D1* play substantial roles in regulating root development.

### Opposite regulation effects of *Rht-D1b* between above-ground and below-ground tissues contribute to a higher root**-**shoot ratio

The opposite regulation effect of *Rht-D1b* in coleoptile and root systems and the dwarfing effect of *Rht-B1b* on coleoptile prompt us to suppose that these two GR alleles function in regulating the root-shoot ratio (RSR) and that the *Rht-D1b* has a larger effect. The fresh weight of shoots and roots at seven DAGs was measured to address this hypothesis. Firstly, the RSR significantly increased in both *Rht-B1aRht-D1b* and *Rht-B1bRht-D1a* genotypes compared with that of *Rht-B1aRht-D1a* genotypes in the natural population, and the *Rht-B1aRht-D1b* genotypes showed a larger increase (Fig. 9a). Then, the *Rht-D1b* was both significantly associated with RSR in GWAS with all genotypes and the subset genotypes (*Rht-B1aRht-D1a* and *Rht-B1aRht-D1b*) as the input, and account for 18.07% and 21.94% of the phenotypic variation explained (PVE), respectively (Fig. 9b and Table S19). The *Rht-B1b* was only significantly associated with RSR with the subset genotypes (*Rht-B1aRht-D1a* and *Rht-B1bRht-D1a*) as the input, accounting for 16.68% of the PVE. Consistently, both the NIL lines Paragon*^B1aD1b^* and Paragon*^B1bD1a^* showed significantly increased RSR than that of Paragon*^B1aD1a^*, and the Paragon*^B1aD1b^* showed a larger increase than Paragon*^B1bD1a^* (Fig. 9c).

**Figure 9.**
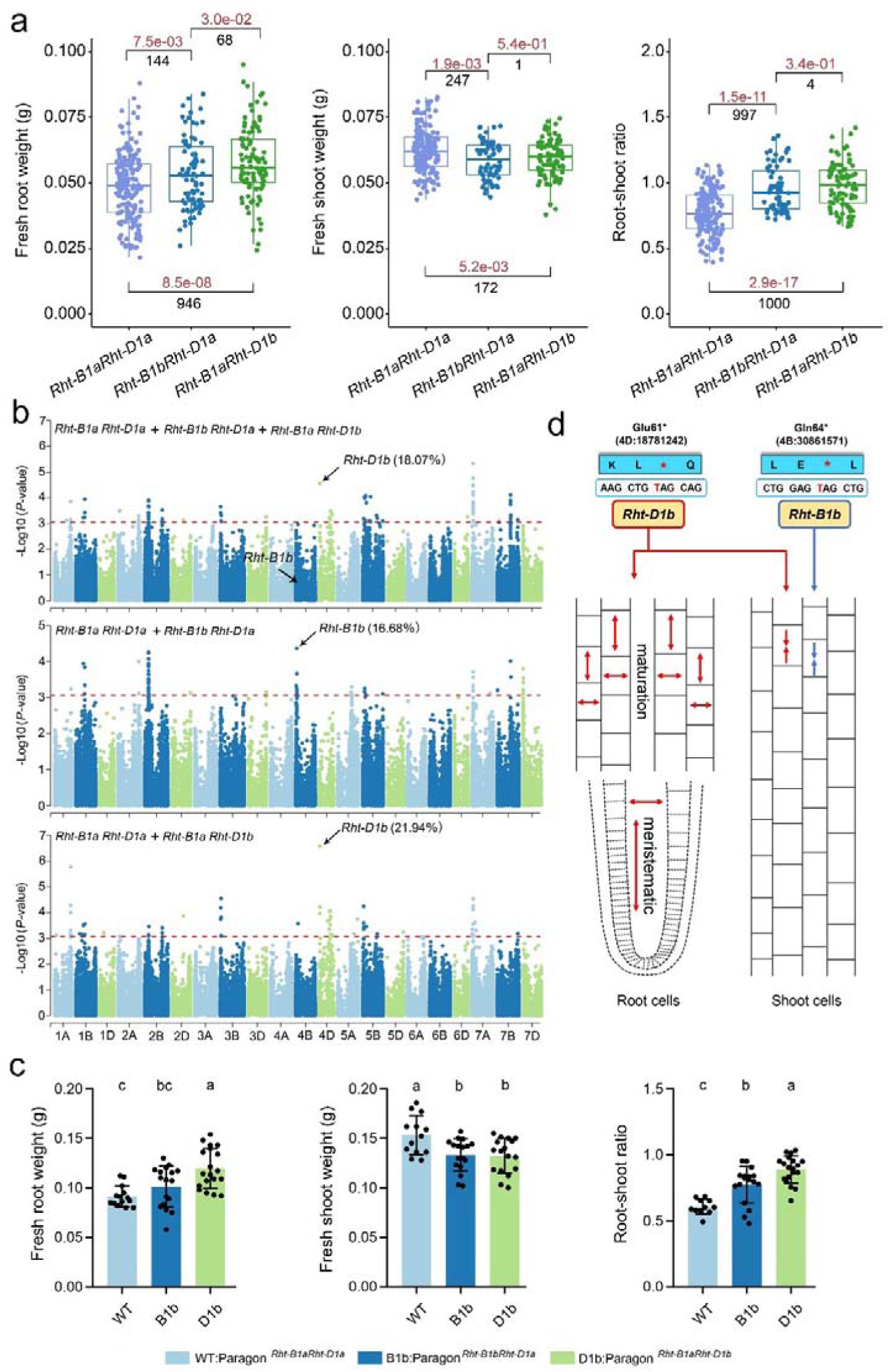
*Rht-D1b* increases the root-shoot ratio much more than *Rht-B1b*. (a) The comparisons of root weight, shoot weight and root-shoot ratio among *Rht-B1aRht-D1a* (n = 171), *Rht-B1bRht-D1a* (n = 78) and *Rht-B1aRht-D1b* (n = 94) genotypes. Red numbers indicate the *P-*value from Student’s *t*-tests, and “NS” represents the insignificance difference at *P* > 0.01 level. Black numbers represent the observed numbers of significant or insignificant differences at *P* < 0.01 or *P* > 0.01 in the 1,000 permutation tests. (b) GWAS results of the root-shoot ratio. The genotypes used for each GWAS analysis were labelled. The number indicates the phenotypic variance explained by *Rht-D1b* or *Rht-B1b*. (c) Statistical data of the near-isogenic lines’ root weight, shoot weight and root-shoot ratio. Data are means ± SD, and different letters (a, b, and c) on the top indicate significant differences (*P* < 0.05) with multiple comparisons by least significant difference (LSD) test. 10-20 individuals were investigated for each line. (d) Model diagram of *Rht-D1b* and *Rht-B1b* regulating root and coleoptile development.

Collectedly, the introductions of *Rht-D1b* conferred distinct responses between above- and under-ground tissues, increasing the cell length and width in roots while inhibiting the cell length in the coleoptile, conferring a larger root system and a higher RSR at the seedling stage (Fig. 9d).

## Discussion

With the release of the reference sequence of Chinese Spring and genome sequence assemblies in the pangenome project^2,43,44^, the scientific studies of bread wheat, including the identifications and characterizations of genes regulating agricultural traits, have stepped into the fast lane. However, the larger linkage disequilibrium (LD) hindered the determination of causal genes in the candidate genomic intervals, and the hexaploid nature challenged the functional characterizations of targeted genes. Our study showed that large-scale population transcriptome sequencing data is powerful for addressing these questions in bread wheat, i.e. TWAS facilitate the determination of causal genes, and the detections of genes co-expressed with the targeted gene could provide clues for the functional characterizations. More importantly, we developed a series of methods that put the above strategies into force in wheat studies^38,42^, providing references for a broader range of researchers.

The wheat GR alleles, *Rht-B1b* and *Rht-D1b*, have dramatically increased grain yields and they are now present in >70% of current commercial wheat cultivars worldwide^21,22^. The improved lodging-resistant and harvest index conferred by these two alleles were regarded as the main reasons for their effects on yield boosting^41,45^. Several studies have investigated whether *Rht-B1b* and *Rht-D1b* also modulated the root development and finally contributed to yield boosting but did not draw consistent conclusions^23,24^, which is possible because the used wheat genotypes, genetic backgrounds and cultivation substrates are different among these studies. Our result that *Rht-D1b* significantly increased root systems of wheat seedlings was acquired with 406 wheat accessions and was further validated in different genetic backgrounds and cultivation substrates, providing strong evidence and conclusions for this field. On the other side, the PRL and TRL were used in most of the previous studies, however the extent to which *Rht-D1b* modulated these two traits was much weaker than that of RD and the related RS and RD in our GWAS analysis, which may also account for the previously inconclusive results. Despite either an increased root length or root number being considered a common way to enlarge the root system, the increased root diameter enlightens a new direction for root improvement, which may reconcile the competition between the root length and root number to form a large root system^46^.

Interestingly, we found that *Rht-D1b* has stronger effects on GA signalling inhibition (Fig. S15) and a broader range of regulated genes (Fig. 5) than *Rht-B1b* in root development, consistent with the previous reports that some root phenotypes differed between *Rht-B1b* and *Rht-D1b* NILs^23,24^. Given that *Rht-B1b* and *Rht-D1b* are homoeolog copies, have comparable mutations and have similar effects on reducing plant height, investigations of why they have divergent functions in the regulations of root development would be intriguing. It was worth noting that, in our previous study^47^, *Rht-D1b* was observed to be preferentially adopted in the Yellow and Huai River valleys wheat production region of China, implying the *Rht-D1b* and the larger root system are beneficial for the adaptations of *Rht-D1b* cultivars to the drought and cold winter ecology conditions in this production region. Furthermore, the functional divergence in regulating root development between these two alleles may be linked to their differential worldwide geographical distributions^48-50^.

In addition to the beneficial effects, *Rht-B1b* and *Rht-D1b* also produced some adverse effects for modern wheat production, such as reduced nitrogen use efficiency and carbon fixation, decreased biomass, spike size, grain weight, coleoptile length and seedling leaf area^45,51,52^. Therefore, many alternative dwarfing genes that do not disrupt GA signalling have been cloned and are considered solutions for raising future dwarf wheat cultivars^51-53^. Here, based on the beneficial effects of *Rht-D1b* on root development, we argue that the advantages of these two GR alleles are not understood clearly, and their replacement with other dwarfing genes would be more complex than we thought. On the other side, a deep understanding of molecular mechanisms underlying the beneficial and adverse effects of these two alleles and the relative genetic manipulation is also an important approach to overcoming the adverse effects, such as improving nitrogen use efficiency in rice GR varieties^45,54^.

In the model plants and crops, it is well established that the GA-deficient mutants, such as *ga1-3* in *Arabidopsis*, *sd1* in rice, and *dwarf1* in maize, are often associated with reduced root length and decreased lateral root formation, and the altered GA sensitivity mutants, such as *slr1* in rice, show varying root lengths depending on whether they exhibit enhanced or reduced sensitivity to GA^55-62^. It is worth noting that some GA-related mutants may also have thicker and stronger roots, which might compensate for reduced elongation and help in maintaining stability and nutrient absorption^63-66^. Consistently, our results showed that *Rht-D1b* significantly increased the root diameters of wheat seedlings, which mostly accounts for the observed increased RS and RD in modern wheat cultivars. However, exogenous GA negatively affected root length in wheat in the previous reports^67^ and our results, which seems contrary to that in the model plants. These pieces of evidence illustrated the multifaceted role of GA in shaping root system architecture, affecting not just elongation but also the overall root structure, and suggested the species-dependent roles of GA in regulating root development.

Furthermore, in *Arabidopsis*, root growth was only retarded when *gai*, a non-degradable, mutant DELLA protein, was specifically expressed in endodermal cells^60,61^, suggesting the abundance of GA and DELLA proteins are root tissue-specific for their genetic effects. Significantly, the cases of *Rht-B1b* and *Rht-D1b* are more complex. Both these two alleles introduce premature stop codons in the open reading frames, which might be expected to prevent translation into a functional protein, but they are gain-of-function alleles that encode functional repressors of GA signalling^41^. Peng *et al*. proposed that TR at an alternative ATG codon closely following the mutant stop codons generate N-terminally truncated proteins lacking the DELLA domain, conferring their gain-of-function phenotypes, and the molecular evidence supporting this hypothesis was reported recently^18,19,41^. In our results, the N-terminal truncated CDS of *Rht-D1* had the function of enlarging the root system, in contrast, the N-terminal truncated CDS of *Rht-B1* did not have this function, suggesting the TR may underly the genetic effects of *Rht-D1b.* Next, the efficiency and root tissues of the TR for *Rht-B1b* and *Rht-D1b* would be critical for understanding the differential effects of these two GR alleles on root development.

## Methods

### Plant materials and Root development trait measurement

The 406 bread wheat (*Triticum aestivum* L.) accessions used in this study included 87 landraces (https://wgb.cimmyt.org/gringlobal/), 43 historical cultivars (the cultivars released before 1970), 190 modern cultivars (the cultivars released after 1970), and 60 breeding lines (Table S1). Similar-sized seeds were selected for each accession and sterilized with 2% sodium hypochlorite for 10 minutes. Then, they were rinsed 3-5 times with distilled water. Twenty sterilized seeds of each accession were incubated at 4 L in darkness for 2 days to break dormancy. The germinated seeds of each accession were placed on damp filter papers in the germination box (12×12×5 cm) and cultivated under 22 L/16 L day/night (50% relative air humidity) and 16 h light (2000 Lux)/8 h dark cycles. After 14 days, seven highly similar robust seedlings were collected for each accession. One seedling root was immediately frozen in liquid N2 for the subsequent RNA isolation. Other six samples were used for phenotyping total root length, root surface, root volume, and root diameter with the Wseen LA-S image system (Hangzhou Wseen Testing Technology Co. LTD) and manually measuring the number of roots and the primary root length. Six independent replications were performed for each accession, and 36 root sampling data and six independent samples were produced for RNA preparation. The dynamic root development measurements were performed at 4 and 7 DAG seedlings using the same procedure as above.

The root-related traits of the near-isogenic lines (*Rht-D1b* and *Rht-B1b* in cv. Paragon background) and three F_2_ populations (Yannong22 × Qingfeng1, Ningmai12 × Zhongluo08-3 and MY72 × Changwei18) were evaluated with the same processes as that used for the phenotyping of the 406 accessions.

### RNA sequencing and Reads mapping

Six independent samples from six replications (one root sample per replication) were equally bulked for total RNA extraction and the RNA-seq libraries generation in collaboration with Frasergen Bioinformatics Co., Ltd. (Wuhan, China). Briefly, the paired-end sequencing libraries with an insert size of ∼250 bp were sequenced on the Illumina HiSeq X Ten platform. Using Trimmomatic (v0.33)^68^, raw reads were trimmed to remove sequencing adapters, and reads with low-quality bases were culled. Reads were then aligned to the bread wheat reference genome (IWGSC RefSeq v1.0) using the STAR software (v2.4.2a)^69^ in the 2-pass mapping mode. Reads were also mapped to the transcriptome sequence (IWGSC RefSeq v1.1 annotation) with BWA (v0.7.17)^70^. Uniquely mapping reads that had the same mapping loci on both genome and transcriptome and the proper mapping distances with the relatively paired reads (< 63,068 bp based on the largest intron length in the whole genome) were selected to reduce mapping errors and used for the subsequential SNP calling.

### SNP calling and Annotation

A two-step procedure^25^ was employed with a few modifications. In detail, the raw SNPs were called with a population SNP-calling manner referring to the best practices of GATK (v4.0.2.0)^71^. The “MarkDuplicates” module (Picard-2.17.11) was used to mark the duplication alignment, the “SplitNCigarReads” module (GATK v4.0.2.0) was used to remove the sequences spanning intronic regions, and the “HaplotypeCaller” module (GATK v4.0.2.0) was used to generate individual variants for each sample. SNPs were filtered with the following parameters: mapping quality ≥ 40, SNP quality ≥ 30, genotype quality for each accession ≥ 20, QD (SNP quality/reads depth) ≥ 2; each SNP was more than 5 bp away from an InDel; for homozygous genotypes, the supporting reads ≥ 5 for each accession; for heterozygous genotypes, the supporting reads for both the reference and alternative alleles ≥ 3 for each accession. SNPs that did not meet these parameters were termed missing. To further exclude possible false polymorphic sites caused by intrinsic mapping errors mainly resulting from the homologous on the reference genome and mapping bias inherent to the mapping algorithm, SNPs identified in the accession of wheat cv. Chinese Spring, used to generate reference genome sequences, were defined as error-prone SNP sets. Any SNP that matched the error-prone set was culled. The SNPs that met the filtering criteria were termed high-quality and annotated using the SnpEff (v4.3)^72^. This SNP calling pipeline can be downloaded from https://github.com/biozhp/Population_RNA-seq.

### SNP validation

To evaluate and improve the reproducibility of the pipeline and the accuracy of the final SNPs, a modulated transcriptome was generated with the reference transcriptome (IWGSC RefSeq v1.1) and passed to SNP calling with the above pipeline. Only 387 alternative alleles (0.1%) were detected, suggesting a low false-positive rate of our pipeline. Then, three accessions (s542, s543 and s544) with two biological replications were selected from the population for RNA-seq library construction and SNP identification, showing an average concordant rate of 98.85% between the two-replication RNA samples, which indicates that the SNP calling pipeline was highly reproducible (Table S20). At last, four accessions (K147, K95, s542 and s543) were selected to genotype with wheat 660K SNP arrays, and the consistent ratio of the overlapped SNPs was > 99% (Table S2). Thus, these results proved the accuracy and reliability of our SNP data. In turn, these validations provide key clues to improve the parameters in the pipeline.

### SNP imputation

Beagle (v5.4)^73^ was used to impute missing genotypes. All heterozygous genotypes were masked as missing genotypes before imputation. To get the optimal imputation accuracy and filling rate, 28,257 sites were randomly masked as missing, with the missing rate varied from 10 to 90%, and were subsequently imputed using default parameters. The imputation accuracy was measured by determining the proportion of correctly inferred genotypes to the total masked genotypes, finding that the imputation accuracy was more than 99% when the missing rate was < 0.8 (Table S21). SNPs with a missing rate < 0.8 were imputed by combining imputation accuracies and filling rates.

### Population structure analysis

RAxML (v8.2.12)^74^ was used to construct a maximum-likelihood tree using imputed SNPs of MAF > 0.2, and the command “-f a -m GTRGAMMA -p 12346 -x 12346 -# 100”. The phylogenetic tree was visualized in iTOL (v6)^75^. Imputed SNPs with MAF > 0.05 were used to quantify and infer population structures in ADMIXTURE (v1.3.0)^76^. The parameters K = 2-10 were used to identify different groups, and finally, K = 4 was used in subsequent analysis.

### Quantification of gene expression

Reads were aligned to high and low confidence transcripts (IWGSC RefSeq v1.1 annotation) with the default parameters in Kallisto (v0.46.2)^77^. The TPM was summarised to gene expression levels using R package Tximport (v1.14.0)^78^ with the “lengthScaledTPM” option^79^. To investigate the extent of gene expression variation within the population, genes with TPM > 0.5 in at least 5% of accessions (20 accessions) were selected, and the fold change of their TPM at 95th percentiles to that of the 5th percentiles (TPM > 0.5) in the whole population was calculated.

### Transcriptome-Wide Association Study (TWAS)

The TWAS, aiming to calculate the association between the gene expression variation and the phenotype variation among the population, is usually performed with the software developed for genome-wide association study (GWAS) in plants by transforming the continuous variable (gene expression levels) into a discrete variable (“0” and “1”). However, this transformation results in the loss of part of the gene expression information. Here, we used the mixed linear model, the fundamental model in GWAS, to exploit the continuous gene expression variations and exclude the inference of population structure and family relatedness with the lmerTest (v3.1.3) package in R (v 4.2.2) as follows:

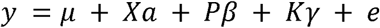

In this equation, *y* represents the values of the investigated phenotype, μ represents the overall mean, Χ is a continuous variable and represents the gene expression values. The *P* and *K* are the PCA and relative kinship matrix generated with GAPIT (v3.1) using the parameters “PCA.total=3”. The top three principal components were used to build the *P* matrix for population-structure correction. The *K* matrix was used to correct the potential familial relatedness. *Xα* and *Pβ* were used as fixed effects, and Κγ was used as random effects, respectively. The genes with false discovery rate (FDR) adjusted *P-*value < 0.05 were considered candidates significantly associated with the investigated phenotypes. Only the genes with expression levels higher than 0.5 (TPM > 0.5) in more than 95% of accessions were used for the TWAS analysis.

### Genome-wide association analyses

The GWAS analysis with the mixed linear model (PCA + kinship) implemented in GAPIT (v3.1)^80^ was applied to the imputed SNPs with MAF > 0.05. The PCA and kinship matrix were generated using GAPIT with the default parameters to correct the population structure and familial relatedness, respectively. A Bonferroni-corrected threshold probability based on individual tests was calculated to correct for multiple comparisons using 0.05/N, where N is the effective SNP number calculated with the Genetic type 1 Error Calculator (v0.2). The significant *P-*value thresholds for root surface and volume were 4.29 × 10^-5^. Those for root-shoot ratio were 4.46 × 10^-5^, 4.52 × 10^-5^, and 4.36 × 10^-5^ for the whole population and the sub-populations removing either *Rht-B1b* or *Rht-D1b* genotypes, respectively. The kernel weight was treated as fixed effects in a linear model to adjust the data of root phenotypes with the “lm” function in R (v 3.6.1)^81^, and the corrected root phenotype data was further applied to the genetic effect analysis and GWAS analysis.

### Identification of SNPs associated with gene expression levels

The genes with expression levels more than 0.5 (TPM > 0.5) in more than 95% of accessions were selected to identify SNPs associated with gene expression levels. Firstly, the expression values among the population for each gene were transformed to a normal distribution with the “qqnorm” function in R (v 3.6.1). Then, the hidden and confounding factors contributing to the expression variability were estimated with Bayesian factor analysis implemented in the PEER (v1.0). Then, the linear mixed models implemented in the GEMMA (v0.98.3) and MatrixEQTL (v2.3) packages that consider the population structure, genetic relatedness and the estimated confounding factors were used for association analysis between the SNPs with MAF > 0.05 (genotypes) and the normalised expression profile of a given gene (expression trait). The significantly associated SNPs (*P*-value < 6.367e-8, determined by 0.01/Total SNP number) generated by the GEMMA and the MatrixEQTL were intersected to improve reliability and accuracy.

### GO enrichment analysis

GO annotations of high-confidence genes were downloaded from the Ensembl Plants Genes 53 database, and GO annotations of low-confidence genes were obtained with eggNOG-mapper (v2)^82^. GO enrichment analysis was performed using the R package clusterProfiler (v3.18.1) with the “enricher” function^83^. The *P*-values were adjusted for multiple tests using Benjamini and Hochberg’s procedure.

### Permutations of the phenotyping comparison

The 1,000 permutation tests were conducted in phenotyping comparison to minimise false positive errors. In each permutation test, 60% of accessions were randomly sampled from each of the two compared groups, and the significance level of the differences was calculated using Student’s *t*-tests. The number of significant differences (*P*-value < 0.01) in the 1,000 permutation tests was recorded.

### Phytohormone treatment

The NIL lines were treated with 0.2 µM, 1 µM and 2 µM gibberellic acid (GA3), 0.5 µM, and 5 µM PAC. Control plants were treated with mock solutions of distilled water containing ethanol. The sterilized seeds (highly similar-sized seeds were selected for each accession) were incubated in the dark at 4 °C for two days, then cultivated in a germinating box for one day, then treated with GA3 and PAC until 14 DAGs. For each accession, three independent replications were conducted for each treatment, and at least six plants were used for each replication.

### Over-expression transgenic vector construction and plant transformation

The full-length and N-terminal truncated coding sequence of *TaRht-B1* (from wheat cv. Ningmai13) and *TaRht-D1* (from wheat cv. Kenong199) was cloned and transferred into the pMWB111 vector with *TaRht-B1* native promoter (1866 bp from wheat cv. Ningmai13) and *TaRht-D1* native promoter (1876 bp from wheat cv. Kenong199), respectively. They were then transformed into wheat cv. Fielder using *Agrobacterium tumefaciens*-mediated transformation^84^. All transgenic lines were confirmed by PCR and sequencing. Similar-sized seeds from transgenic lines and wild type were selected for phenotyping to reduce the interference of grain weights on root development.

### Cytological observation of roots and shoots

Confocal laser scanning microscopy (CLSM) was used for cytological observations of roots and shoots. CLSM samples were prepared according to the previous report^85^. Samples were harvested and fixed in 4% glutaraldehyde (in 12.5 mM cacodylate, pH 6.9), desiccated in vacuo for 30 min, after which they were incubated overnight in a fixative at room temperature. After fixation, tissues were dehydrated through different gradients of ethanol absolute (50%, 60%, 70%, 85%, 95% and 100% ethanol absolute) for 30 min per step. Then, tissues were clarified with 2:1 (v/v) benzyl benzoate: benzyl alcohol for a minimum of 1 h. Samples were observed under a confocal microscope coupled to a 488-nm argon laser (Olympus, Fluoview1200, Tokyo, Japan). ImageJ (v1.48) software was used to measure the length and width of the cell.

## Data availability

The raw RNA-Seq data were deposited in the Sequence Read Archive (https://www.ncbi.nlm.nih.gov/sra) under accession numbers PRJNA838764. Genotypic, transcriptomic, and phenotypic data used in this analysis are publicly available from our database (http://resource.iwheat.net/WGPD/). The analysis code can be downloaded from https://github.com/biozhp/Population_RNA-seq/.

## Acknowledgements

This work was supported by the Biological Breeding-National Science and Technology Major Project (No. 2023ZD0402403 to Xiaoming Wang and No. 2023ZD0402401 to Shengbao Xu) and the National Natural Science Foundation of China (31571756 and 31870298) to Shengbao Xu. We thank Weining Song, Xinhong Chen, and Hude Mao for helping with germplasm collection and the support of NWAFU’s High-Performance Computing.

## Author contributions

S.X. and Z.K. conceived the project and designed the research. Z.K., S.X., D.H. and W.J. collected wheat accession. X.W., P.Z., X.G., Z.L. and L.H. performed the sample preparation, RNA sequencing and phenotypic data collection. X.W., P.Z. and Y.L. performed the data analysis. X.S. and W.H. generated the transgenic lines. J.S. and C.U. generated the NIL lines. W.H., M.C., X.L. (Xueting Liu), and X.L. (Xiangjun Lai) measured the root-related traits of NIL lines, segregation populations and transgenic lines. X.G. and J.W. performed the cytological observations. X.W. and S.X. organised the data and wrote the manuscript. C.U. and J.X. provided critical comments to improve the manuscript. All authors discussed the results, commented on the manuscript, and approved the final manuscript.

---

**Figure S1.**
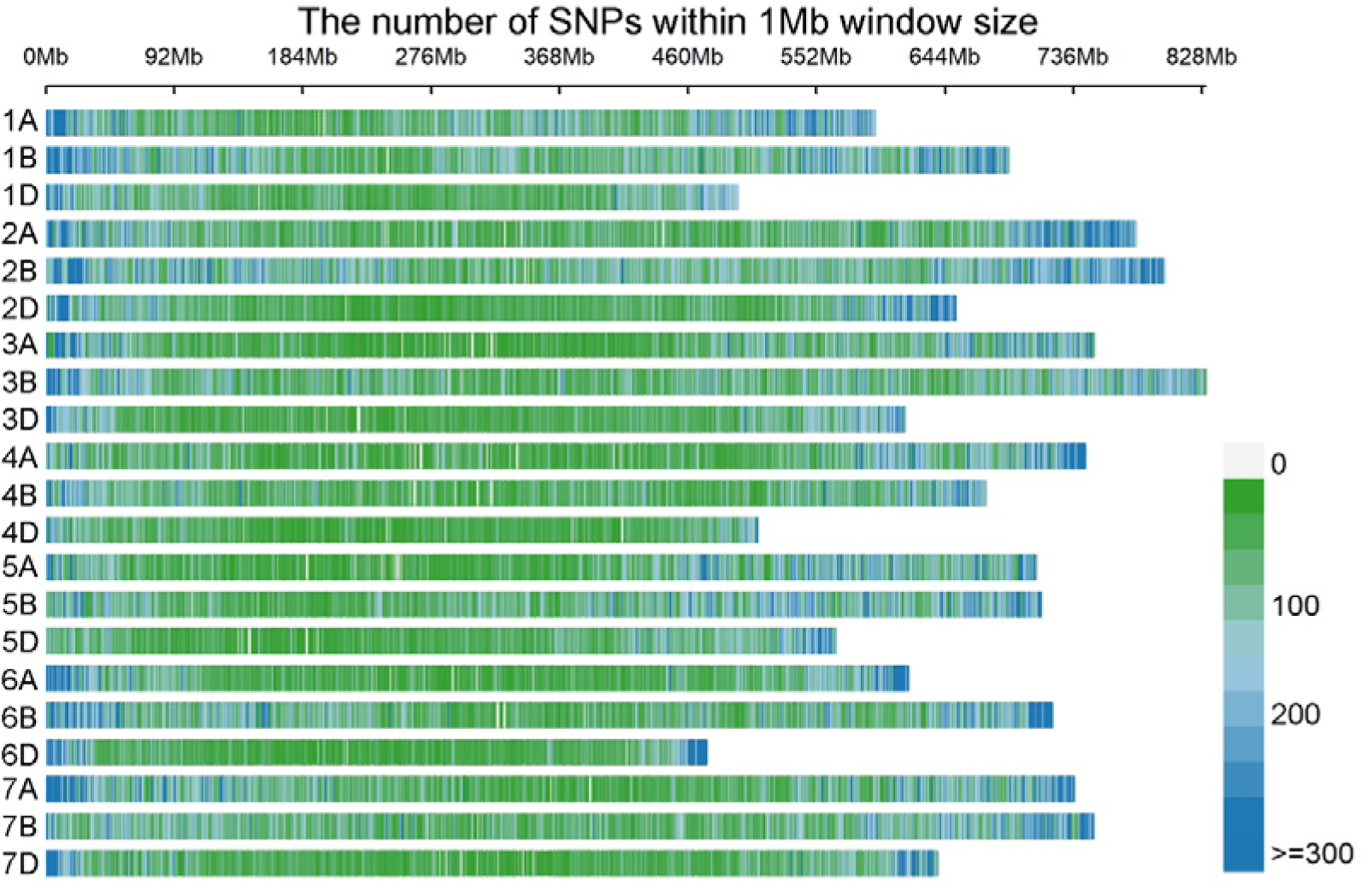
The distribution of the identified SNPs. The colour indicates the number of SNPs within 1LJMb window size.

**Figure S2.**
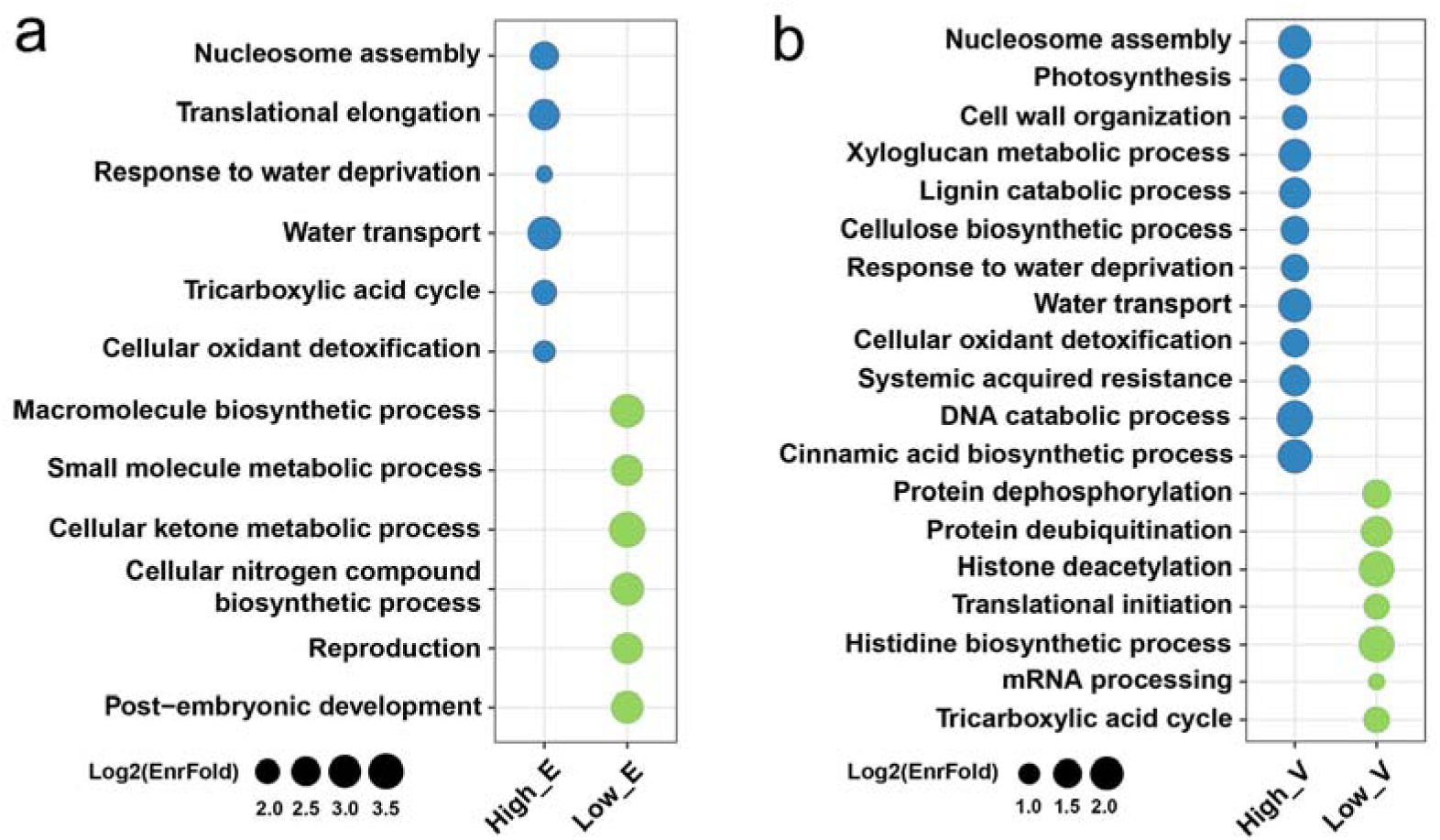
GO enrichment of the genes with highly and lowly expressed (a) or varied (b) genes. (a) The top 10% of genes with the highest expression (High_E) and lowest expression (Low_E) in the population transcriptome were selected for this analysis. (b) The top 10% of genes with the highest variation expressions (High_V) and lowest variation expressions (Low_V) were selected for this analysis.

**Figure S3.**
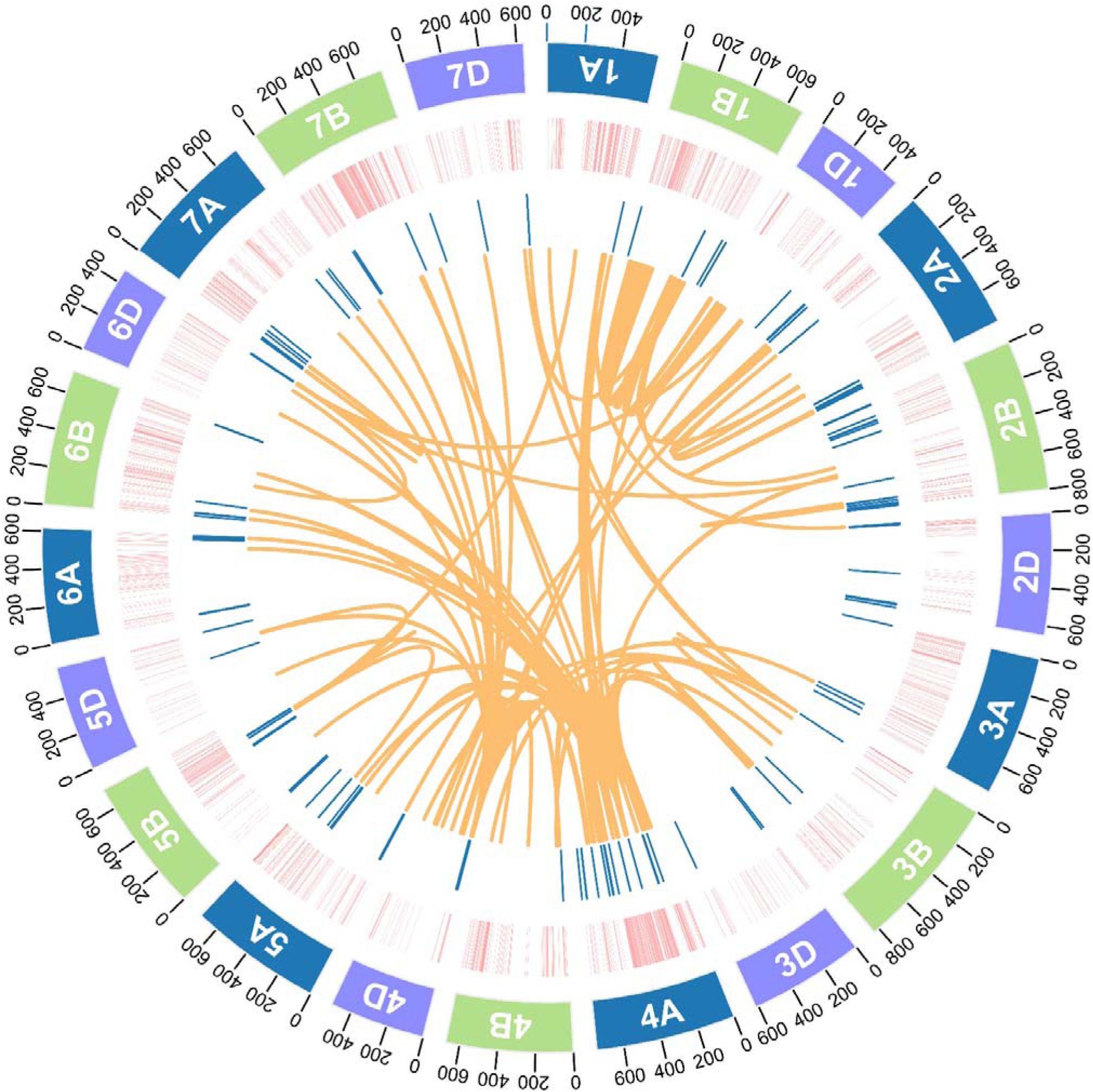
Circos diagram showing the selection sweeps, GWAS results and TWAS results. The lays from outer to inner represent the chromosomes, the selection sweep during wheat breeding, and the SNPs located in the selection sweeps and were significantly associated with root-related traits (MC-LA-SNPs). The lines in the centre represent the link between MC-LA-SNPs and the genes whose expression levels were significantly associated with root-related traits.

**Figure S4.**
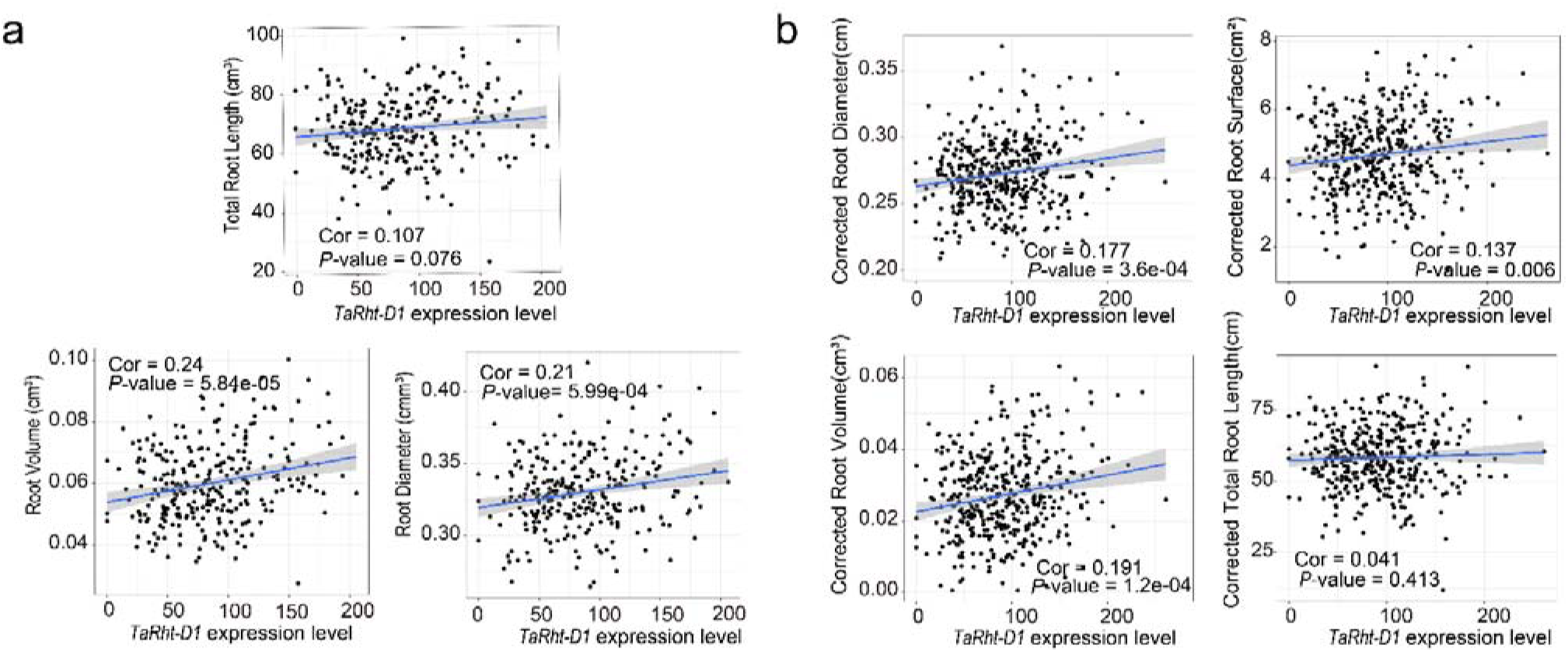
Associations between the expression levels of *TaRht-D1* and the original (a) and corrected (b) root-related traits. In (b), the phenotypes were corrected with the general linear model to exclude the effects of kernel weight on root development.

**Figure S5.**
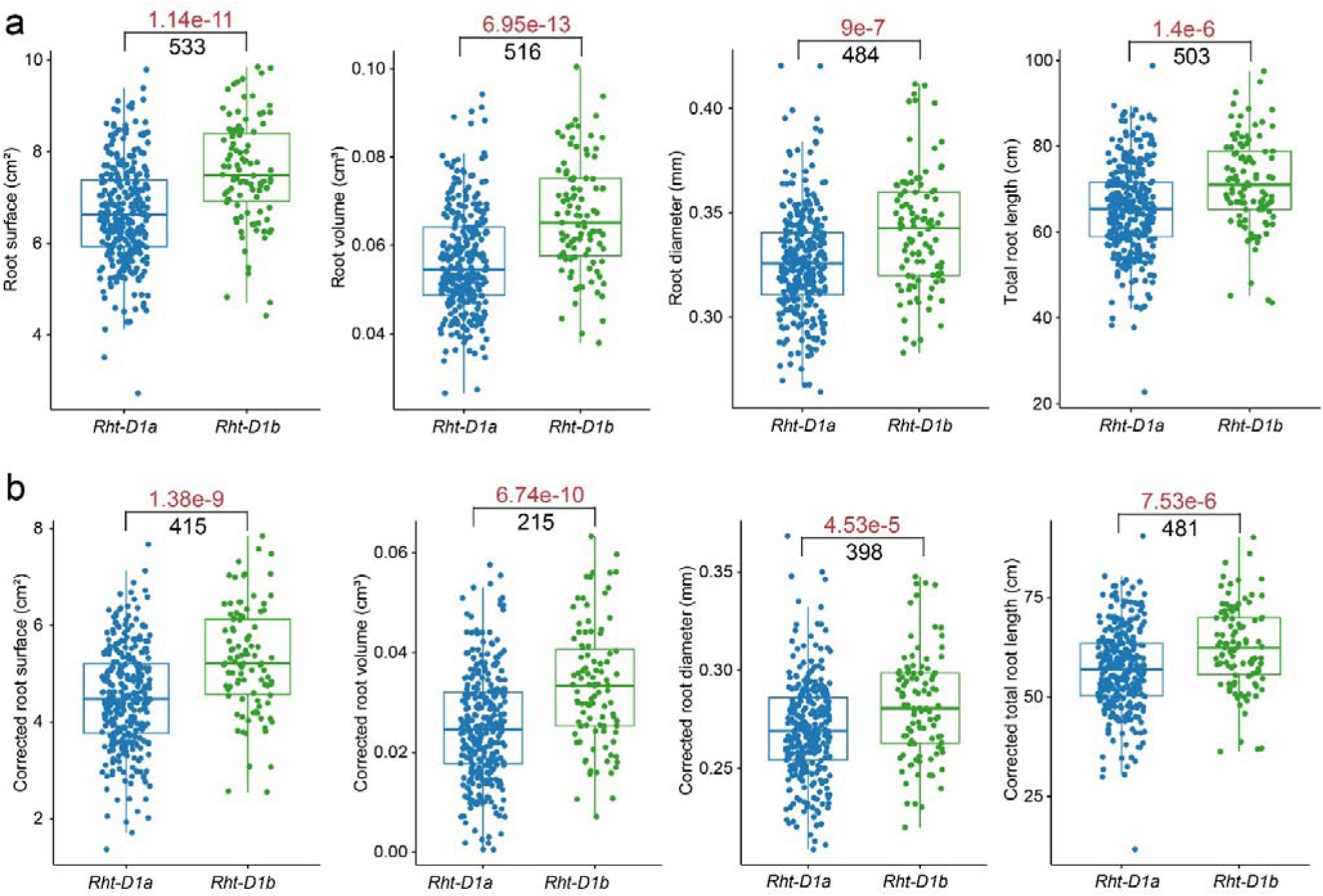
The comparisons of the root-related traits between *RhtD1a* (n= 307) and *RhtD1b* (n= 99) genotypes. (a) Comparisons with the original phenotypes. (b) The phenotypes were corrected with the general linear model to exclude the effects of kernel weight on root development. Data are means ± SD. Red numbers indicate the *P-*value from the Student’s *t*-tests. Black numbers indicate the observed number with significant differences at *P* < 0.01 level in the 1,000 permutation tests.

**Figure S6.**
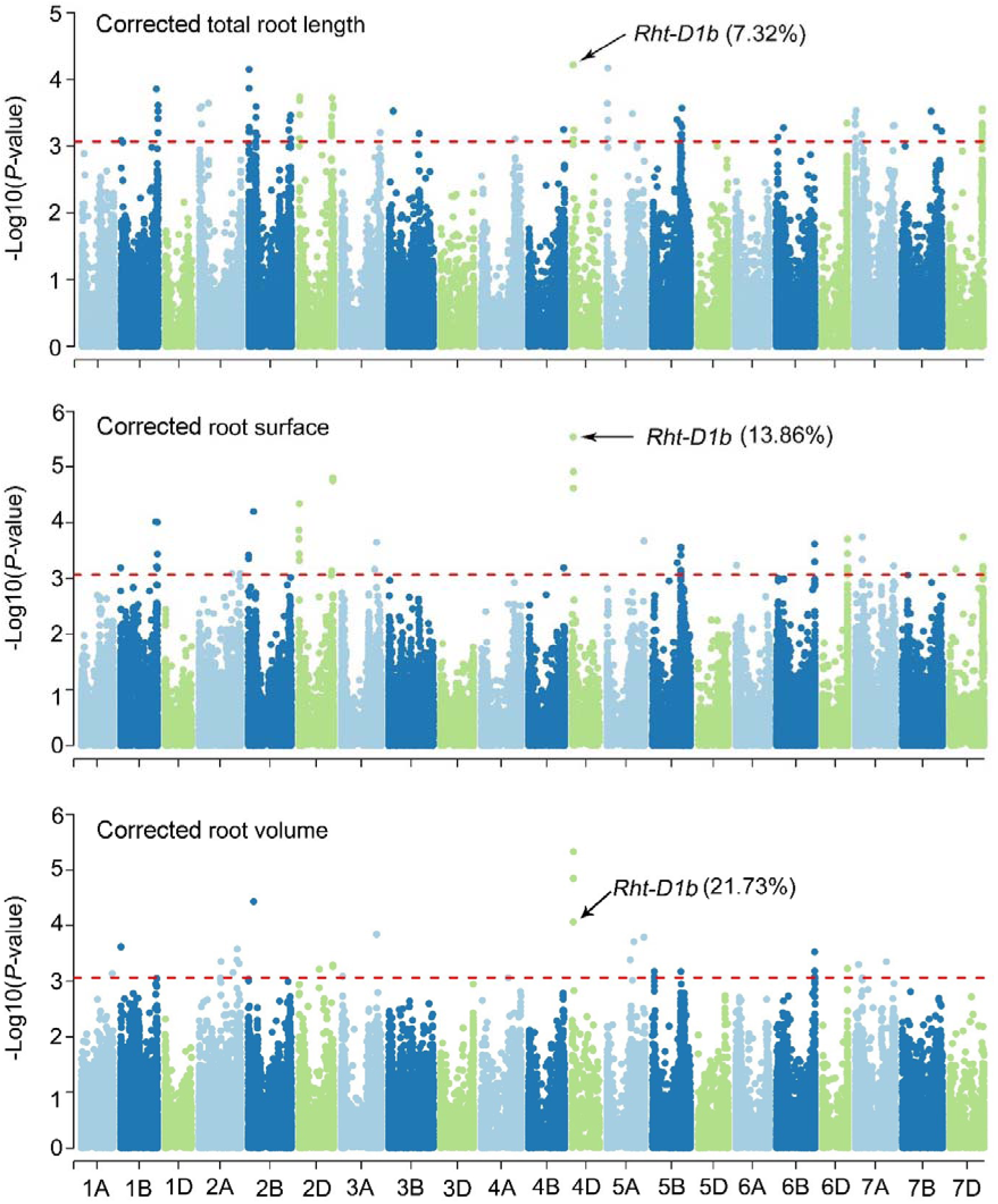
GWAS results of the corrected root-related traits. The phenotypes were corrected with the general linear model to exclude the effects of kernel weight on root development. The genotypes used here were the same as those in Fig. 4a. The red line in each plot represents the significant *P*-value threshold calculated with Bonferroni-corrected threshold probability. The numbers indicate the phenotypic variance explained by *Rht-D1b*.

**Figure S7.**
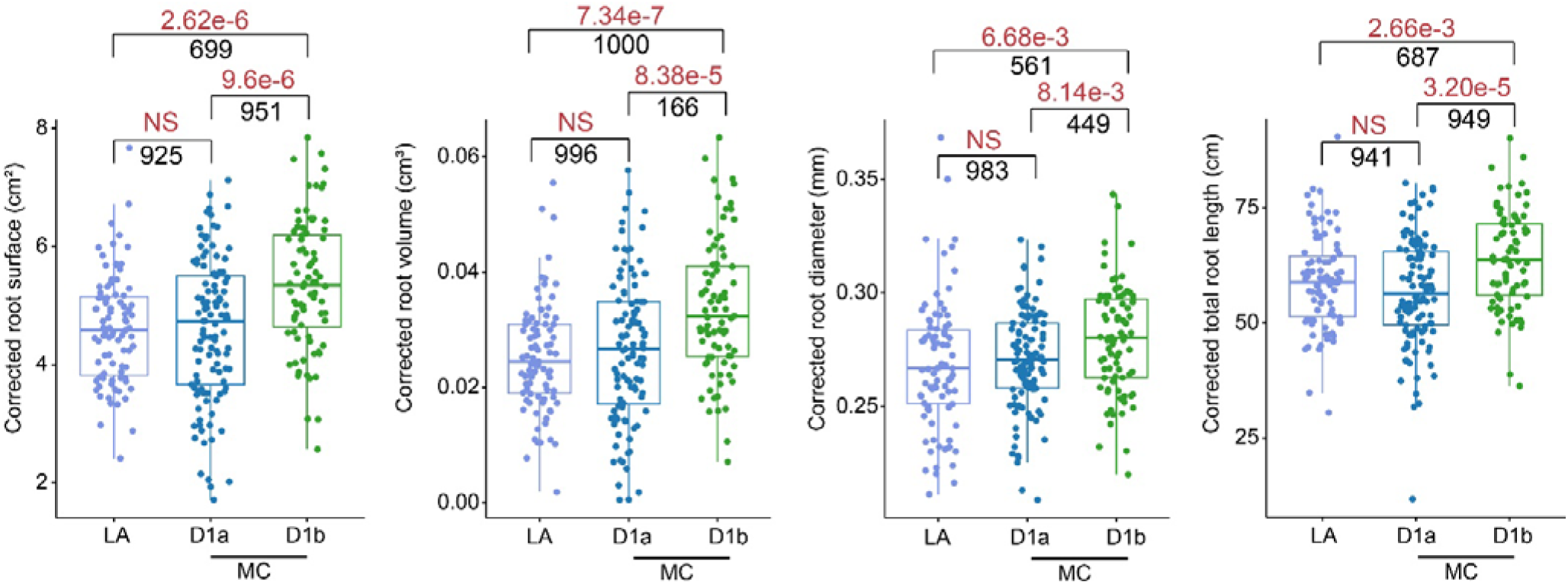
Comparisons of corrected root phenotypes between LA (n = 87) and MC (n = 190) groups. The MC accessions were divided into *D1a* (n = 110) and *D1b* (n = 80) genotypes based on the GR allele *Rht-D1b*. Red numbers indicate the *P-*values from the Student’s *t*-test, and “NS” represents insignificance at the *P*-value > 0.01 level. Black numbers indicate the observed number with significant differences at *P*-value < 0.01 level or insignificant differences at *P*-value > 0.01 level (for the “NS”) in the 1,000 permutation tests.

**Figure S8.**
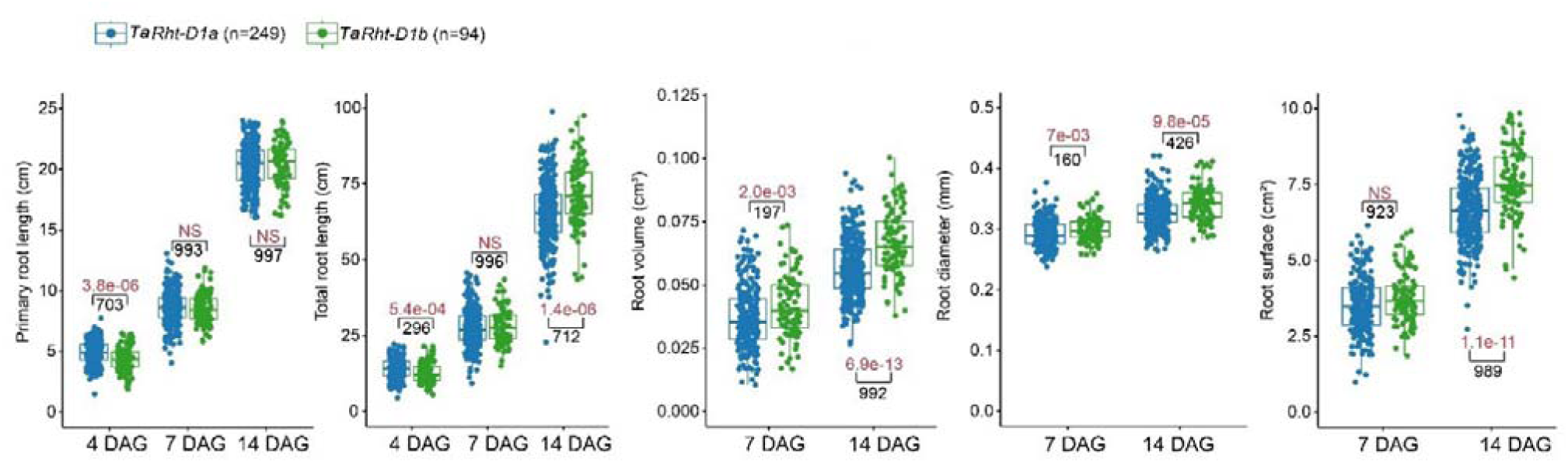
The comparisons of root-related traits between *Rht-D1a* and *Rht-D1b* genotypes at 4, 7, and 14 DAGs (days after germination). Red numbers indicate the *P-*values from the Student’s *t*-test, and “NS” represents insignificance at the *P*-value > 0.01 level. Black numbers indicate the observed number with significant differences at *P*-value < 0.01 or insignificant differences at *P*-value > 0.01 level (for the “NS”) in the 1,000 permutation tests.

**Figure S9.**
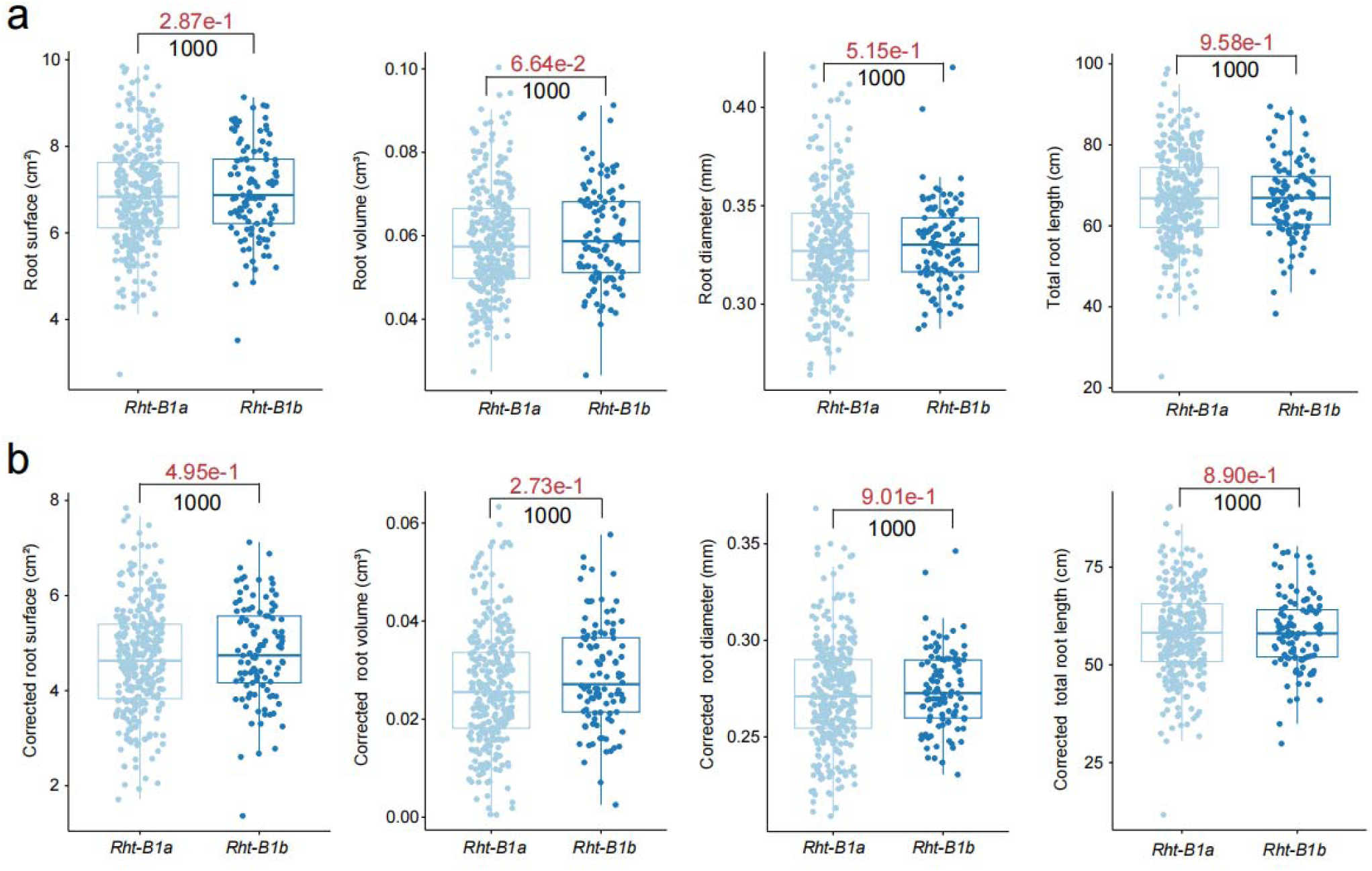
The comparisons of original (a) and corrected (b) root-related traits between *Rht-B1a* (n = 298) and *Rht-B1b* (n = 108) genotypes. Red numbers indicate the *P-*value from the Student’s *t*-tests. Black numbers indicate the observed number with insignificant differences at *P* > 0.01 in the 1,000 permutation tests.

**Figure S10.**
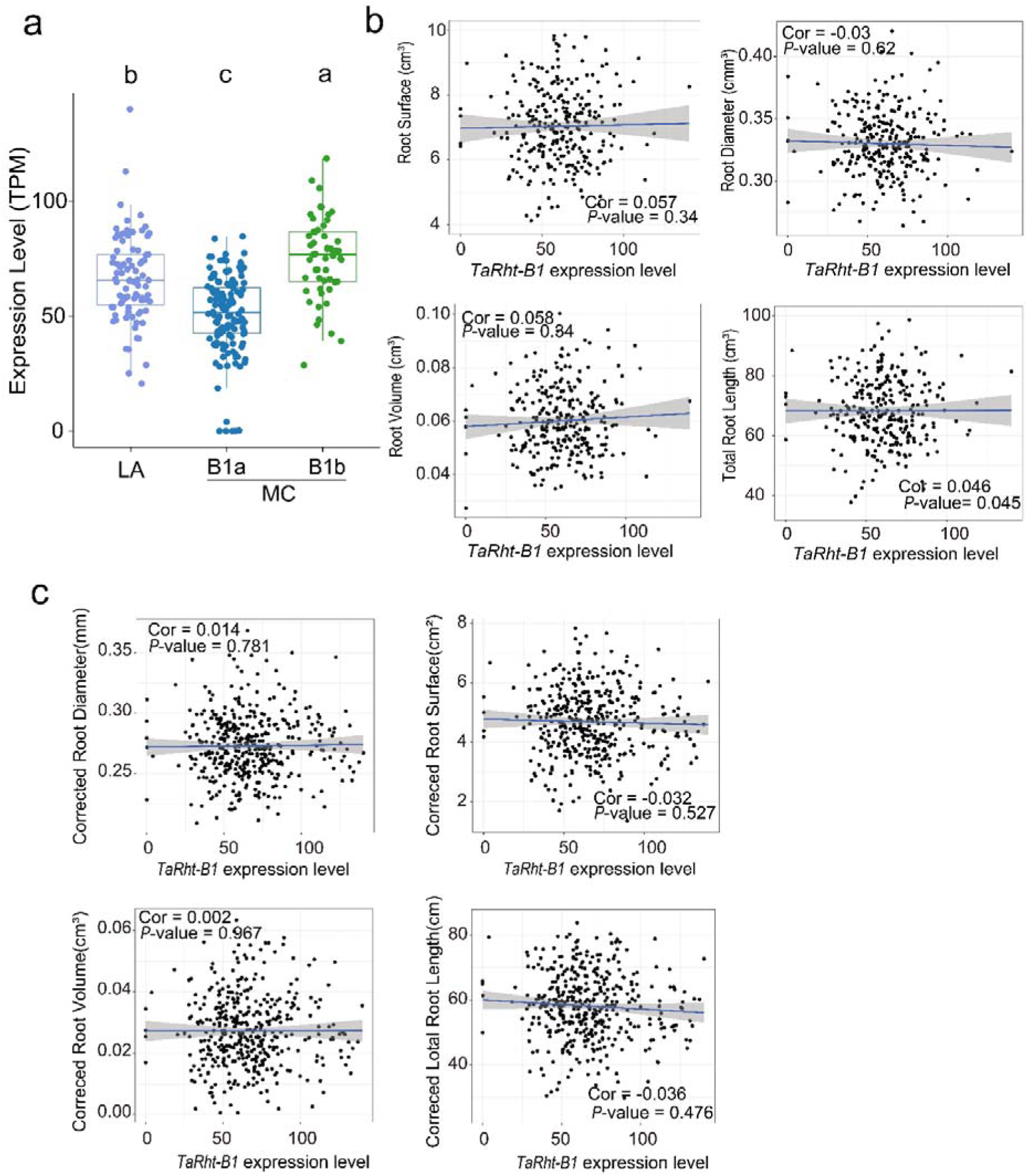
Associations between the expression levels of *TaRht-B1* and the original (b) and corrected (c) root-related traits. **(a)** The expression levels of *TaRht-D1*. The MC accessions were divided into *B1a* (n = 110) and *B1b.* Different letters on the top indicate significant differences (*P* < 0.05) calculated with multiple comparisons by least significant difference (LSD) test. In (c), the phenotypes were corrected with the general linear model to exclude the effects of kernel weight on root development.

**Figure S11.**
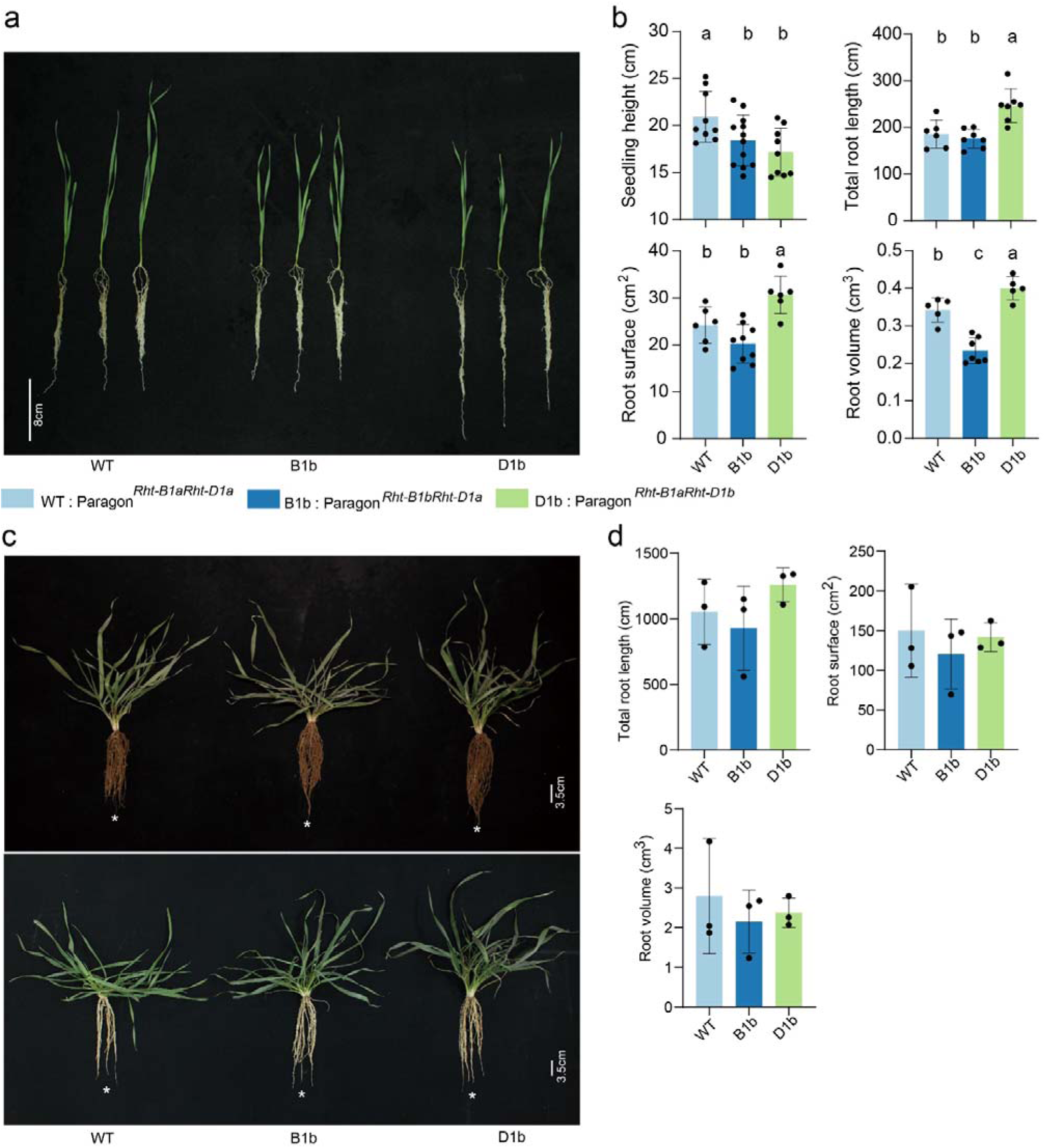
The genetic effects of *Rht-B1b* and *Rht-D1b* on root development. (a-b) Phenotype (a) and statistic data (b) of the Near-isogenic lines cultivated with sand substrate. (c-d) Phenotype (c) and statistic data (d) of the Near-isogenic lines cultivated in the field. Data are means ± SD, and different letters on the top indicate significant differences (*P* < 0.05) calculated with multiple comparisons by least significant difference (LSD) test.

**Figure S12.**
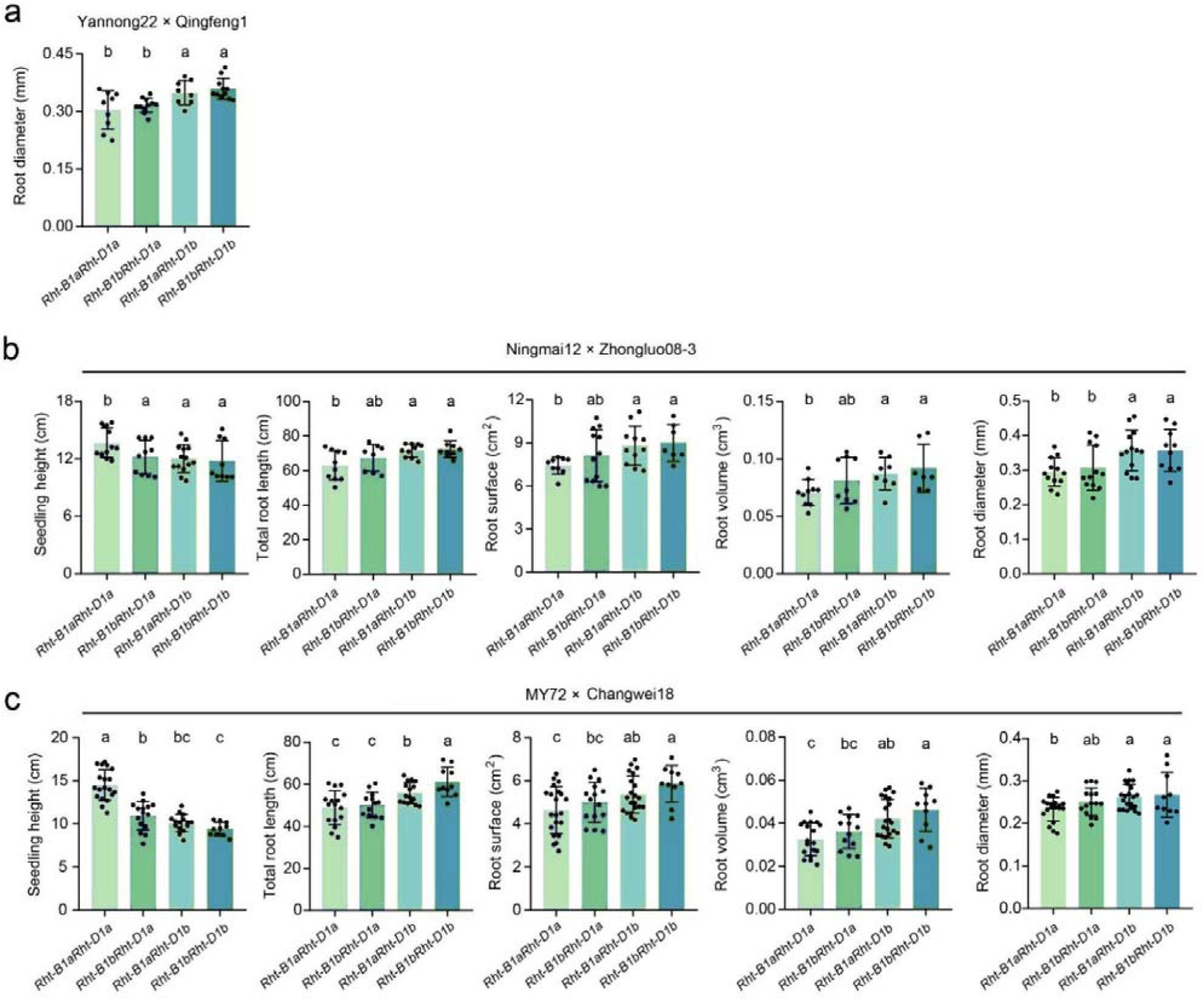
The comparison of seedling height and root-related traits among different genotypes in three F_2:3_ segregation populations. The parents are (a) Yannong22 (*Rht-B1bRht-D1a*) × Qingfeng1 (*Rht-B1aRht-D1b*), (b) Ningmai12 (*Rht-B1bRht-D1a*) × Zhongluo08-3 (*Rht-B1aRht-D1b*) and (c) MY72 (*Rht-B1bRht-D1a*) × Changwei18 (*Rht-B1aRht-D1b*), respectively. Data are means ± SD, and different letters on the top indicate significant differences (*P* < 0.05) with multiple comparisons by least significant difference (LSD) test.

**Figure S13.**
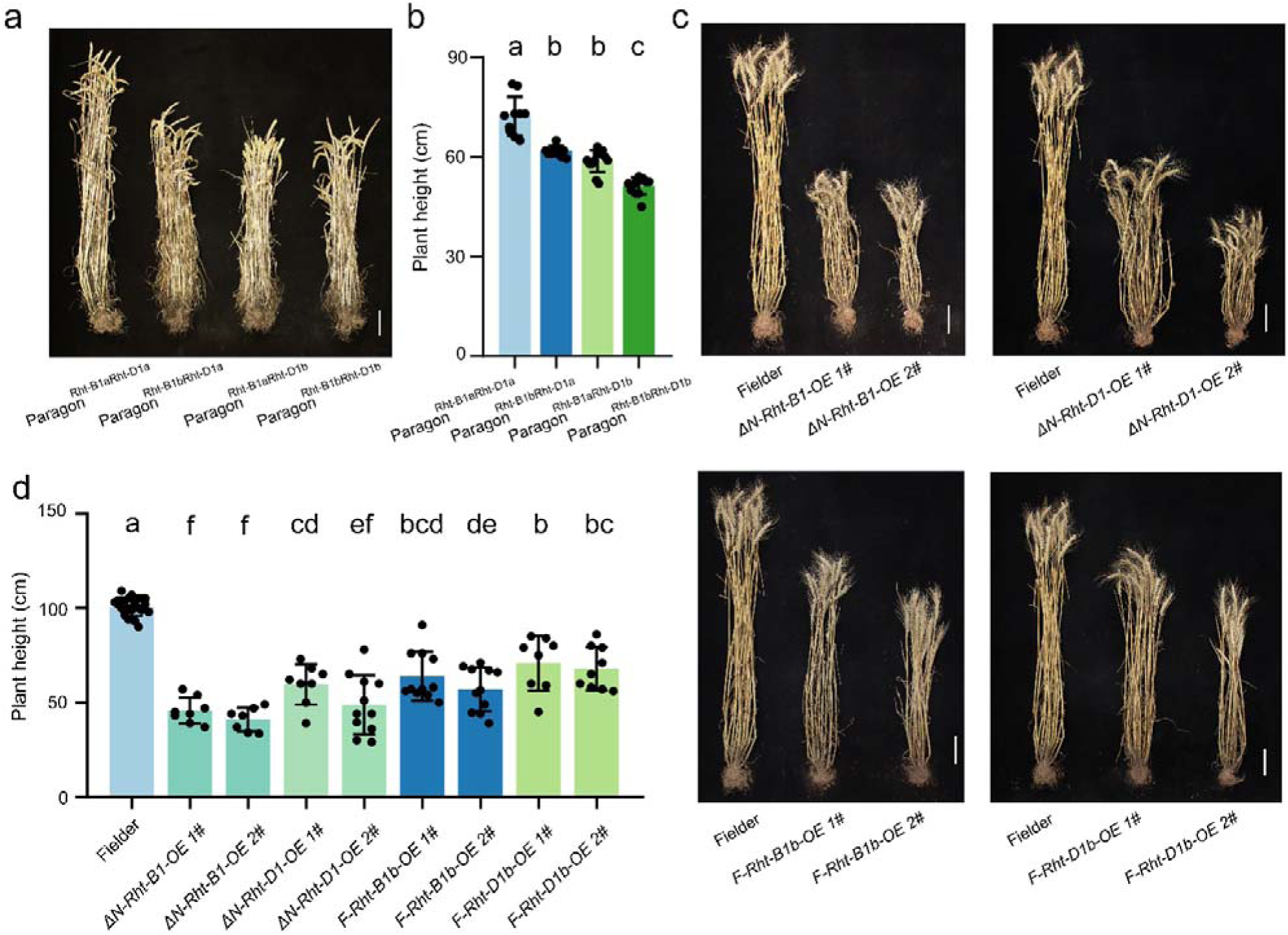
Plant height of the *Rht-B1*/*Rht-D1* transgenic lines and the near-isogenic lines at the mature stage. (a-b) Phenotype (a) and statistical data (b) of plant height of the near-isogenic lines. (c-d) Phenotype (c) and statistical data (d) of plant height in the wild-type, *Rht-B1* and *Rht-D1* transgenic lines. Data are means ± SD, and different letters on the top (b/d) indicate significant differences (*P* < 0.05) between different genotypes with multiple comparisons by least significant difference (LSD) test. Fielder: wild-type; *F-Rht-B1b-OE* 1/2#, full CDS of *TaRht-B1* transgenic lines; *F-Rht-D1b-OE* 1/2 #: full CDS of *TaRht-D1* transgenic lines. Δ*N-Rht-B1-OE* 1/2#, N-terminal truncated CDS of *TaRht-B1* transgenic lines; Δ*N-Rht-D1-OE* 1/2 #: N-terminal truncated CDS of *TaRht-D1* transgenic lines. Bar = 10 cm.

**Figure S14.**
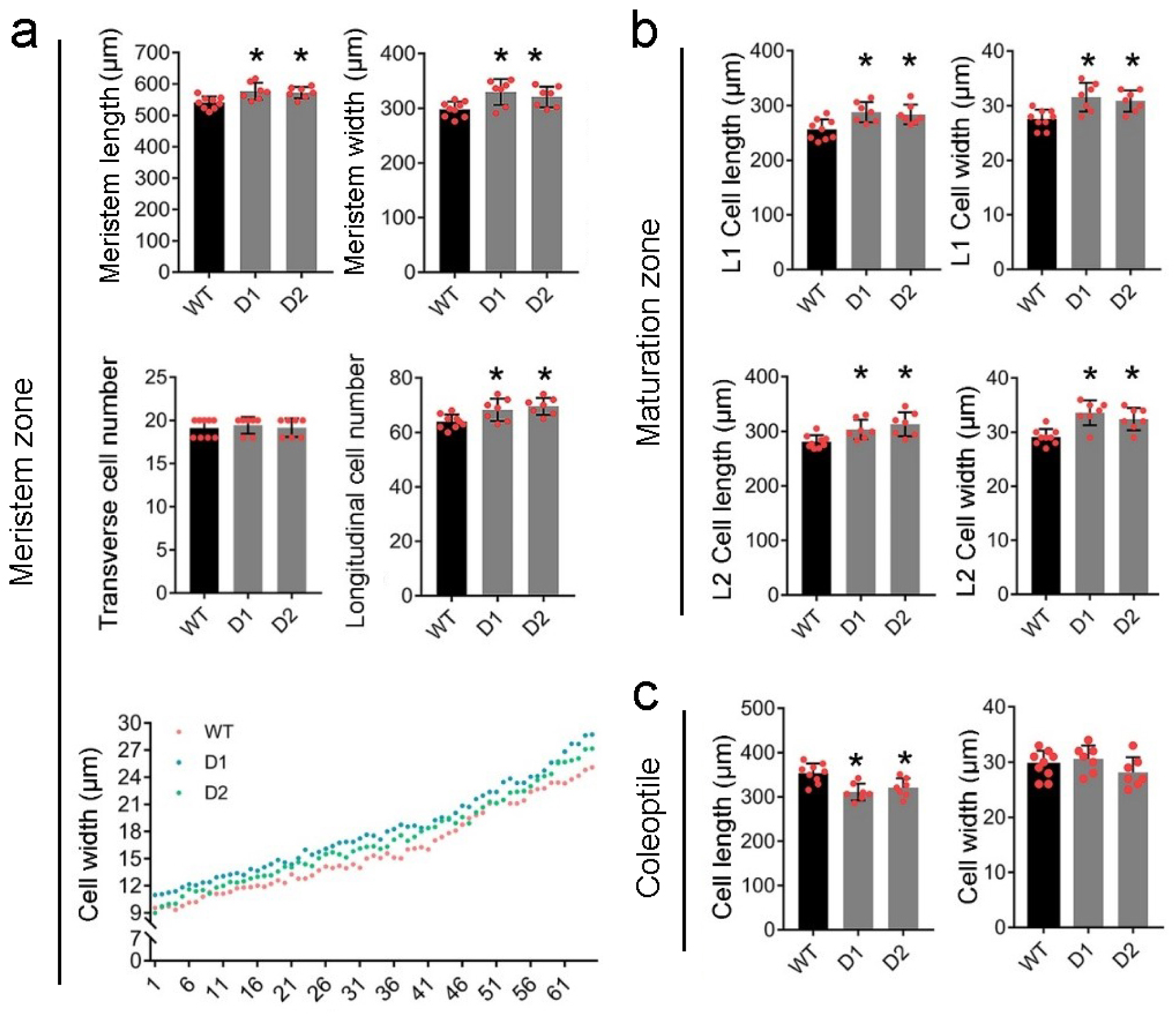
Cytological observations of the *TaRht-D1*’s transgenic lines. (a) Comparisons of root meristem zone of transgenic lines to WT. The longitudinal cell length and number were measured from the cell layer indicated by the red asterisk in Fig. 7a. (b) Comparisons of root maturation zone of transgenic lines to WT. “L1” and “L2” represent different cell layers indicated by the red and yellow dotted lines in Fig.7b. (c) Comparisons of transgenic line coleoptile to that of WT. WT, wild type (cv. Fielder), the D1 and D2 are transgenic lines of *F-Rht-D1b-OE 1#* and *F-Rht-D1b-OE 2#.* At least seven samples were observed for each line, and at least 30 clear cells in each sample were measured with ImageJ software (v1.48). “*” indicate significant differences (Student’s *t*-test P < 0.05) between the compared lines.

**Figure S15.**
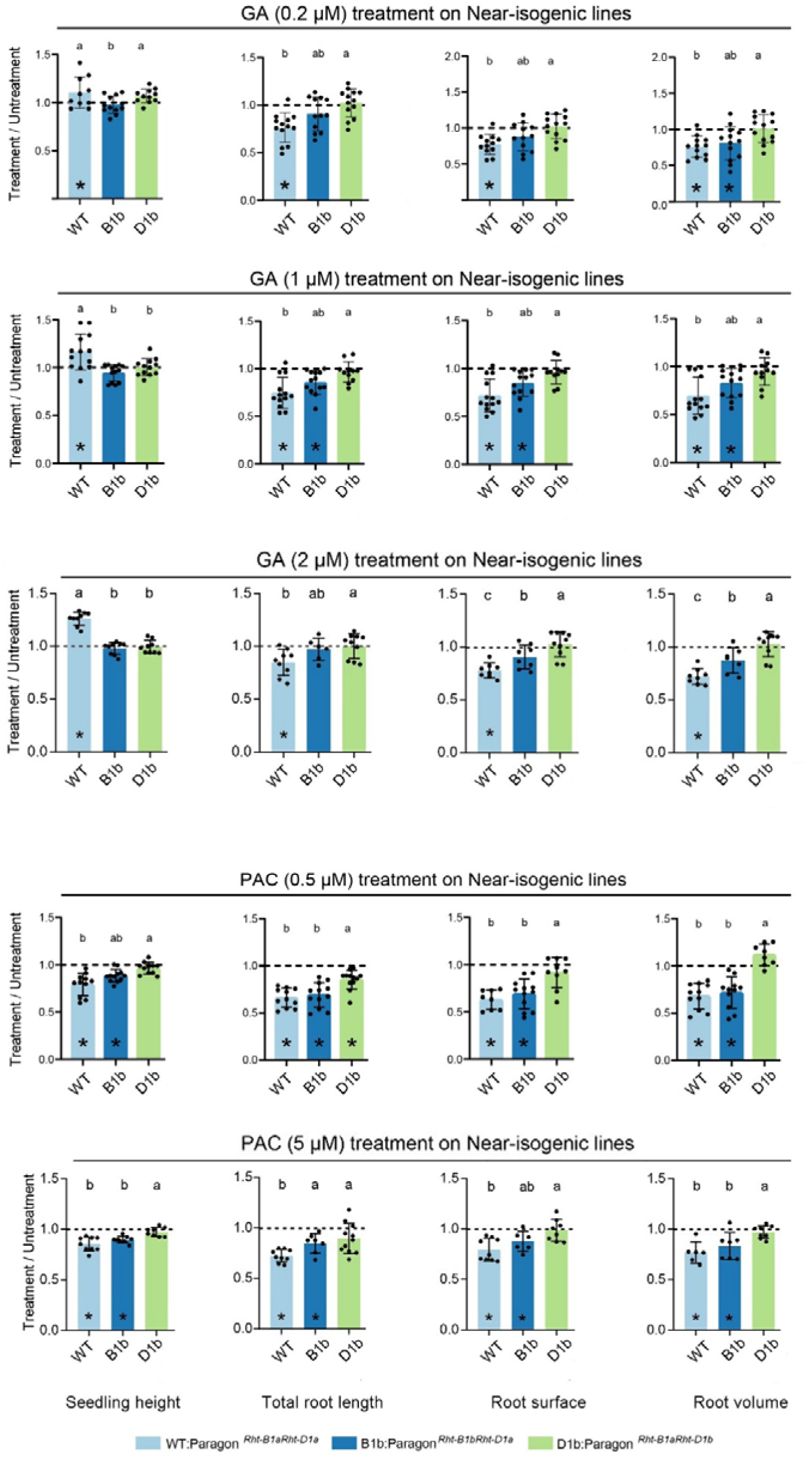
The response of seedling height and root-related traits to GA (gibberellin) and PAC (polybulozole) treatments at different concentrations. The Y-axis represents the ratio of phenotype values of the treated group to that of the untreated group. “*” indicates significant differences (Student’s *t*-test *P* < 0.05) between treated and untreated groups for each NIL line. Different letters on the top indicate significant differences (*P* < 0.05) in the response degree among NIL lines calculated with multiple comparisons by least significant difference (LSD) test.

